# Performing Parentage Analysis for Polysomic Inheritances Based on Allelic Phenotypes

**DOI:** 10.1101/2020.09.15.297812

**Authors:** Kang Huang, Gwendolyn Huber, Kermit Ritland, Derek W. Dunn, Baoguo Li

## Abstract

Polyploidy poses several problems for parentage analysis. We present a new polysomic inheritance model for parentage analysis based on genotypes or allelic phenotypes to solve these problems. The effects of five factors are simultaneously accommodated in this model: (i) double-reduction, (ii) null alleles, (iii) negative amplification, (iv) genotyping errors and (v) self-fertilization. To solve genotyping ambiguity (unknown allele dosage), we developed a new method to establish the likelihood formulas for allelic phenotype data and to simultaneously include the effects of our five chosen factors. We then evaluated and compared the performance of our new method with three established methods by using both simulated data and empirical data from the cultivated blueberry (*Vaccinium corymbosum*). We also developed and compared the performance of two additional estimators to estimate the genotyping error rate and the sample rate. We make our new methods freely available in the software package polygene, at http://github.com/huangkang1987/polygene.

## Introduction

Parentage analysis is a common technique in plant ecology and selective breeding. This technique for identifying parents enables researchers to assess seed dispersal (Ismail *et al*., 2017), pollen dispersal (Bezemer *et al*., 2016), assortative mating (Monthe *et al*., 2017), isolation (Tambarussi *et al*., 2015), current gene flow (Duminil *et al*., 2016), mating systems (Tan *et al*., 2019), reproductive success (Watanabe *et al*., 2018), functional sex (Oddou-Muratorio *et al*., 2018), and to increase genetic gain from selective breeding (Norman *et al*., 2018).

A large proportion of plant species are polyploid, with 24% of all plant taxa exhibiting some form of polysomic inheritance (Barker *et al*., 2016), and at least 47% of angiosperm species having polyploidy in their ancestral lineage (Wood *et al*., 2009). Existing methods of parentage analysis for polyploids use the pseudo-dominant approach (Rodzen *et al*., 2004; Wang and Scribner, 2014) and exclusion approach (Zwart *et al*., 2016). In the pseudo-dominant approach, the polyploid genotypes or the allelic phenotypes are converted into pseudo-dominant phenotypes and use diploid likelihood equations to calculate the likelihood for parentage assignment (Gerber *et al*., 2000), in which each allele at a codominant locus is treated as an independent dominant ‘locus’. This approach enables rapid calculation but is inferior to that based on polysomic inheritance methods because any transformation of data will cause a loss of information and thus a reduction in accuracy (Wang and Scribner, 2014). The exclusion approach excludes the parents based on Mendelian incompatibility. However, due to the high gamete diversity (Pelé *et al*., 2018) and genotyping ambiguity (Huang *et al*., 2014), the exclusion rate is low in polyploid, especially for a parent-offspring pair. Thus, the development of more accurate methods of parentage analysis for polyploids is required.

Several models for polysomic inheritance have been developed, such as double-reduction models (Muller, 1914; Haldane, 1930; Mather, 1935), genotypic frequencies (Fisher, 1943; Geiringer, 1949), and transitional probabilities from a zygote to a gamete (Fisher and Mather, 1943; Field *et al*., 2017). On the basis of these findings, Huang *et al*. (2019) derived the generalized genotypic frequency and gamete frequency for ploidy levels fewer than 12 and derived the generalized transitional probability from a zygote to a gamete for any ploidy level. These models provide a foundation on which to establish a method of parentage analysis for polyploids.

A unique feature of polysomic inheritance is *double-reduction* such that a pair of sister chromatids are segregated into a single gamete (Parisod *et al*., 2010). Double-reduction arises from a combination of three major events during meiosis: (i) the crossing-over between non-sister chromatids, (ii) an appropriate pattern of disjunction, and (iii) the migration of chromosomal segments carrying a pair of sister chromatids to the same gamete (Darlington, 1929; Haldane, 1930). Geneticists have developed several mathematical models to simulate double-reduction: these are the *random chromosome segregation* (RCS) model (Muller, 1914), the *pure random chromatid segregation* (PRCS) model (Haldane, 1930), the *complete equational segregation* (CES) model (Mather, 1935) and the *partial equational segregation* (PES) model (Huang *et al*., 2019). A brief description of each of these models is given in Appendix A.

There are two consequences of double-reduction that will influence parentage analysis: (i) the genotypic frequencies will deviate from expected values, resulting in a bias of the estimated LOD scores and (ii) some unexpected offspring genotypes may be generated (e.g. an offspring genotype *AAEE* is produced from *ABCD* × *EFGH*) along with the true father being excluded. Therefore, the complete array of diverse polyploid offspring genotypes has to be accounted for in order to conduct a comprehensive and accurate paternity analysis (Stift *et al*., 2008, 2010).

There are also several additional problems associated with PCR-based markers that need to accounted for, irrespective of ploidy. One problem is the *genotyping ambiguity* of polyploids (Huang *et al*., 2014), in the sense that the allelic dosage of PCR-based markers cannot be determined. For example, the genotype *AABB* will appear to be identical to *AAAB*. Another problem arises when using microsatellites, which are the genetic markers most frequently used for parentage analysis. Microsatellites can have null alleles (Ravinet *et al*., 2016) that cause both the lack of amplification of null allele homozygotes and the lack of detectability of null allele heterozygotes (Wagner *et al*., 2006). A third problem comes from genotyping errors, which may cause a true parent to be mistakenly excluded due to an observed lack of shared alleles with the offspring (Blouin, 2003). Finally, inbreeding will result in an excess of homozygotes in a population, such as when plants self-fertilize (Ritland, 2002). The genotypic frequencies used for a parentage analysis will thus be affected by any inbreeding.

Here, we extend the disomic inheritance model of Kalinowski *et al*. (2007) to account for polysomic inheritance to enable accurate parentage analysis for polyploids based on genotypes or allelic phenotypes. Our new polysomic inheritance model accommodates the effects of five factors: (i) double-reduction, (ii) null alleles, (iii) negative amplification, (iv) genotyping errors and (v) self-fertilization. To solve the problem of genotyping ambiguity, we develop a new method so as to establish the likelihood formulas for allelic phenotype data, with the effects of our five factors of interest also being included in these formulas. We subsequently use a designated simulated dataset to evaluate and compare the performance of our new method with three other established methods. We also use an empirical microsatellite dataset from the cultivated blueberry (*Vaccinium corymbosum*) to test the performance of all four methods. Moreover, we develop and evaluate two models to estimate the genotyping error rate and the sample rate. We have incorporated our new parentage analysis methods in to the software package polygene, which can be freely downloaded at http://github.com/huangkang1987/polygene.

## Theory and modelling

Here we assume that our parentage analysis model satisfies four assumptions, which are also commonly used for diploid population genetics methods. These four assumptions are: (i) the population is large enough to negate any effects of genetic drift and there is no population subdivision; (ii) the mating is not only random but also independent of both the genetic markers used and the parental mating system, (iii) the distributions of the genotypes are the same for males and females, and reach an equilibrium state (i.e. genotypic frequencies do not change among generations) and (iv) the genetic markers used are autosomal, codominant and unlinked.

The multiset consisting of allele copies within an individual at a locus is called a *genotype*, denoted by 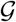 or *G*, in which 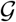 represents an observed genotype and *G* represents a true genotype. For example, {*A, A, A, B*} is a genotype, abbreviated as *AAAB*. The set consisting of alleles within an individual at a locus is called an *allelic phenotype*, or a *phenotype* for short, denoted by 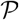. For instance, {*A, B*} is a phenotype, written as *AB* for short.

Our methods are the extensions of Kalinowski *et al*.’s (2007) method. In the following text, we briefly describe the scheme of Kalinowski *et al*.’s (2007) method and its associated diploid model.

### Scheme of simulation-based likelihood approach

The foundations for assigning parentage with confidence by a simulation-based likelihood approach were establish by Marshall *et al*. (1998). There are three typical categories in this approach: (i) identifying the father (or one parent) when the mother (or the other parent) is unknown; (ii) identifying the father (or one parent) when the mother (or the other parent) is known; and (iii) identifying the father and the mother (or parents) jointly. There are two situations in the third category, the first is for dioecious species and the sexes of individuals are recorded (termed sexes known), and the second is for monoecious species or the sexes of individuals are not recorded (termed sexes unknown). The procedures of a parentage analysis are broadly as follows.

For each of the first two categories, two hypotheses are established: the *first hypothesis* is that the alleged father is the true father, denoted by *H*_1_; the *alternative hypothesis* is that the alleged father is not the true father, denoted by *H*_2_. For the third category, ‘father’ needs to be changed to ‘parents’ in both hypotheses.

Given a hypothesis *H*, the *likelihood* is defined as the probability of some observed data given *H*, written as 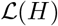. Returning to *H*_1_ and *H*_2_ as described above, we call the natural logarithm of the ratio of 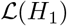 to 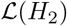 the *LOD score*, or *LOD* as the abbreviation, symbolically 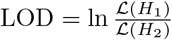. Moreover, if a LOD is positive, it means that *H*_1_ is more likely to be true than *H*_2_. Similarly, a negative LOD means that *H*_2_ is more likely to be true than *H*_1_.

Marshall *et al*. (1998) provided a statistic Δ for resolving paternity, the definition of which is:

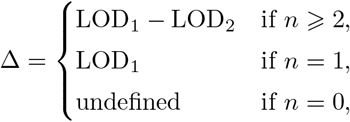

where LOD_1_ and LOD_2_ are respectively the LODs of the most-likely and the next most-likely alleged fathers, and *n* is the number of all alleged fathers. For a practical application, the statistic Δ needs to be singly calculated for each individual offspring. Monte-Carlo simulations are subsequently used to assess the confidence level of Δ. The symbol Δ_0.95_ represents that the threshold of Δ reaches the confidence level 95%, in the sense that if Δ ⩾ Δ_0.95_, the probability that the assigned parent is the true parent is at least 0.95.

The likelihood equations used in Marshall *et al*. (1998) to accommodate genotyping error miscalculate the probability of observing an erroneous genotype. Therefore, we applied the corrected equations in Kalinowski *et al*. (2007) in the following.

### Marshall *et al*.’s (1998) diploid model

Marshall *et al*.’s (1998) diploid model (abbreviated as the Ma-model) accounts for any genotyping errors under the assumption that the genotype frequencies accord with the Hardy-Weinberg equilibrium (HWE). This model consists of some likelihood formulas (listed in the first half of Appendix B) together with the rules and methods for a general parentage analysis.

The likelihood formulas of the Ma-modelare derived by using the transitional probability 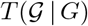 from a true genotype *G* to an observed genotype 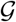, whose expression is

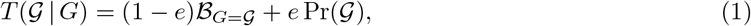

where *e* is the genotyping error rate, 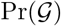 is the frequency of 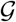, and 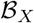 is a binary variable, such that 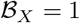 if the expression *X* is true, or 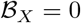 otherwise.

As previously stated, the procedures underlying the Ma-modelto perform a parentage analysis are as follows: (i) calculating 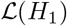 and 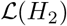, (ii) finding the threshold of Δ, (iii) calculating the LOD and Δ, and (iv) using the values obtained in the previous three steps to assess the confidence level of this parentage analysis.

In the following text, we will use the first category in a parentage analysis as an example to show how to calculate the likelihoods 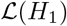 and 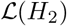 in the Ma-model. The expressions of 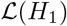 and 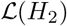 are

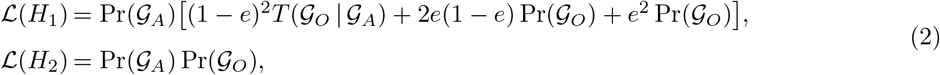

where 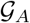 and 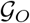 are respectively the observed genotypes of the alleged father and the offspring, 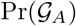 and 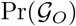 are their frequencies, and 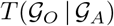 is the transitional probability from 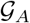 to 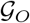.

In the Ma-model, the genotyping error is considered as the replacement of a true genotype with a random genotype according to the genotypic frequencies. Thus the genotyping error does not change the distribution of the genotypes, i.e. 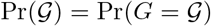. Moreover, Pr(*G*) can be directly calculated from the HWE prediction:

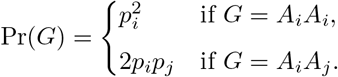

This is because any null alleles, any negative amplification (i.e. amplification failure due to experimental error or a poor DNA quality, rather than a null allelic homozygote) and any inbreeding/selfing are not considered in the Ma-model.

Next, the transitional probability 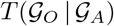 is calculated under the assumptions that 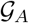 and 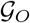 are correctly typed and that the alleged father is the true father, i.e. under the assumptions that 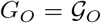 and 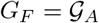, where *G_O_* and *G_F_* are the true genotypes of the offspring and the true father, respectively. Therefore, 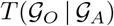 is the same as *T*(*G_O_* | *G_F_*) under these assumptions. Because one allele within *G_O_* is randomly inherited from the parents, and the other is randomly sampled from the population according to the allele frequencies, the transitional probability *T*(*G_O_* | *G_F_*) can be expressed as

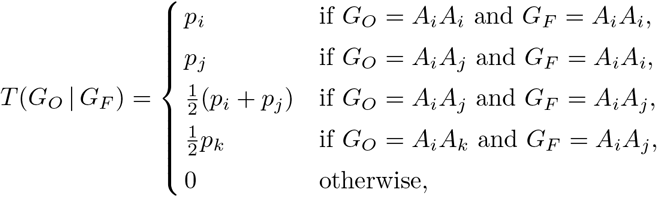

where *A_i_*, *A_j_* and *A_k_*. are distinct identical-by-state alleles, *p_i_, p_j_* and *p_k_* are their frequencies.

Now, we see that the two likelihood formulas in Equation (2) can be used for the actual calculation as long as the values of the genotyping error rate *e* and those frequencies of alleles are given.

For the second and third categories in a parentage analysis, to calculate the transitional probabilities 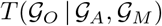 and 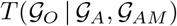 in the likelihood formulas in the Ma-model(see the first half of Appendix B), we need to apply the transitional probability *T*(*G_O_* | *G_F_*, *G_M_*) from a pair of true genotypes of the true parents to a true genotype of the offspring. Because the genotypic frequencies in the Ma-modelaccord with the HWE, according to the Mendelian segregation (i.e. each parent randomly contributes one allele to an offspring genotype), 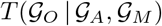 can be calculated by

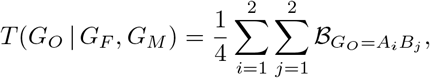

where *A_i_* (or *B_j_*) is an allele within *G_F_* (or *G_M_*).

### Polyploid model

The polysomic inheritance model (abbreviated as the *polyploid model*) presented here is for use with even levels of ploidy, and consists of some likelihood formulas and some additional conditions along with the rules and methods for a general parentage analysis. These additional conditions are: (i) which of the two data types (genotypic and phenotypic) are to be selected, (ii) whether self-fertilization is considered, (iii) whether null alleles and/or negative amplifications are to be considered, and (iv) which of the four double-reduction models, listed in Table S1, is chosen.

As for the Ma-model, our new model accommodates the effect of genotyping errors and the presence of these errors will not change the genotypic and phenotypic frequencies. Moreover, if self-fertilization is considered in our model, its effect will also be incorporated into the likelihood formulas.

For the genotypic data, the likelihood formulas for all three categories in a parentage analysis, under either self-fertilization or not, are given in Appendix B. For polysomic inheritance, the genotypic frequencies 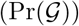 and transitional probabilities (*T*(*G_O_* | *G_F_*) and *T*(*G_O_* | *G_F_*, *G_M_*)) need to be properly adjusted, where the formula of 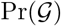 under inbreeding and double-reduction is given in Appendix C (or in Huang *et al*. (2019)), and the formulas of *T*(*G_O_* | *G_F_*) and *T*(*G_O_* | *G_F_*, *G_M_*) are given in Appendix D.

For the phenotypic data, the likelihood formulas for all three categories in a parentage analysis under the condition of either self-fertilization or not are given in Appendix E. In such circumstances, the phenotypic frequencies 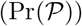 in these formulas are calculated by Equation (A5), and the transitional probabilities 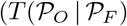 and 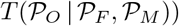 by Equation (3) or (4). To solve the problem of genotyping ambiguity, we develop a new method termed the phenotype method. In this method, the prior probabilities of phenotypes and the transitional probability from a phenotype to another phenotype will be used to establish various likelihood formulas.

### Phenotype method

We begin our discussion with the symbol 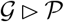, whose meaning is that 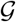 is a genotype determining the phenotype 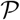, i.e. 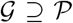 and 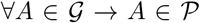, where ⊇ is the inclusion of multisets. If the null alleles (e.g. *A_y_*) are considered, the conditions should be revised to 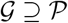 and 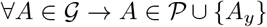. Under the revised conditions, our models will accommodate the effect of null alleles.

The formulas of transitional probabilities 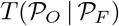 and 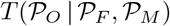 are first established, whose expressions are

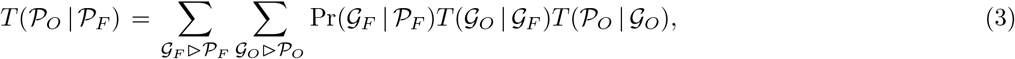

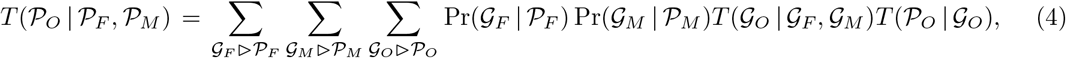

where 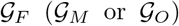 is taken from all genotypes determining 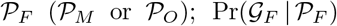 and 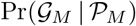 are two posterior probabilities, which can be calculated by the Bayes formula

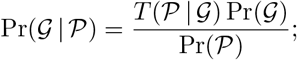

and 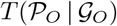 is the transitional probability from 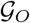 to 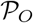, which is calculated by

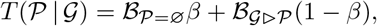

in which *β* is the negative amplification rate, and 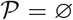 means that 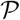 is a negative phenotype (it may be caused by either a null allele homozygote or a negative amplification).

Because each genotype may encounter an amplification failure, the candidate genotypes determining a negative phenotype at a locus are, strictly speaking, all possible genotypes at this locus. This will create a problem for the calculations of the transitional probabilities. This is because there are up to 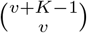 genotypes at a locus, where *v* is the ploidy level and *K* is the number of alleles at this locus. For example, the number of genotypes at an octo-allelic locus for tetrasomic (hexasomic, octosomic or decasomic) inheritance is up to 330 (1716, 6435 or 19448). For this reason, we do not consider the candidate genotypes determining any negative phenotypes. In other words, all negative phenotypes are discarded in the polysomic inheritance model during the analytical process. However, they will still be used in the allele frequency estimation so as to estimate the negative amplification rate *β* and the null allele frequency *p_y_*.

Next, the likelihood formulas for all three categories are established. For example, if self-fertilization is not considered, the likelihoods 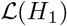 and 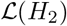 for the first category can be simply obtained by replacing 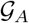 with 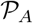 and 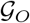 with 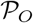 in Equation (2), whose expressions are

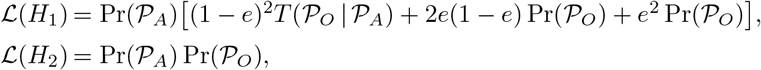

where 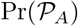 and 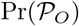 are respectively the frequencies of 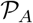 and 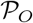, which can be calculated by Equation (A5), and the transitional probability 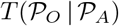 is calculated by replacing 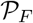 with 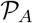 in Equation (3), i.e. 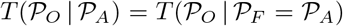. The likelihood formulas for each category under the condition of either self-fertilization or not are given in Appendix E.

### Estimation of genotyping error rate

For a genotypic dataset, it is mathematically impossible to estimate the genotyping error rate *e* without any additional information (e.g. the information of pedigree or replication). We will develop a genotyping error rate estimator based on the pedigree data, including the known parents and the identified parents (at a high confidence level, e.g. 99%). We refer to a parent-offspring pair extracted from the pedigree data as a *reference pair*, and a father-mother-offspring trio as a *reference trio*.

For genotypic data, we assume that the allelic dosage is known so there are no null alleles. For the phenotypic input, all candidate genotypes and their gametes will be extracted, including the genotypes with null alleles, and the pair (or trio) mismatch is identified by whether the parent (or the parents) is able to produce the offspring (see Appendix I for details). Therefore, each mismatch in our models can only be caused by genotyping errors or the false parent(s). Pair mismatches can be used in all three categories, but trio mismatches can only be used in the second and the third categories. In this section, we will use pair mismatches to describe how to estimate the genotyping error rate.

Let *δ* be the probability of observing a pair mismatch in a true parent-offspring pair under the condition that any individual has been erroneously genotyped. In our genotyping error model, *δ* is equal to the exclusion rate for the first category, i.e. the probability that two random genotypes are mismatched. We do not estimate *δ* by simulation or by allele frequencies because those approaches can be influenced by the errors in the estimated parameters. Instead, we directly estimate *δ* from the input genotypes/phenotypes with a Monte-Carlo algorithm, whose procedures are broadly as follows: randomly sample a large number of individual pairs from the input samples with replacement, and then treat each as a parent-offspring pair, and finally calculate the probability that their genotypes/phenotypes at a locus are mismatched, which is used as 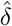 at this locus.

Let *γ* be the probability of observing a pair mismatch in a true parent-offspring pair. Since each mismatch observed in the true parent-offspring pairs can only be caused by the genotyping error, if we denote *E* for 1 – (1 – *e*)^2^, then *γ* = *Eδ*. Noticing that the estimate 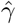 can be calculated from the reference pairs in a single application or in all available applications based on the same dataset, the single-locus estimate 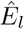 of *E* at the *l^th^* locus can be expressed as 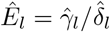.

If we assume that there are *n_rl_* reference pairs at the *l*^th^ locus and that *n_ml_* is the number of pair mismatches in these reference pairs, then *n_ml_* as a random variable obeys the binomial distribution B(*n_rl_*, *γ_l_*), so Var(*n_ml_*) = *n_rl_γ_l_*(1 – *γ_l_*). Because 1 – *γ_l_* is close to one, the variance 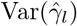 can be approximately expressed as 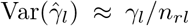. Because 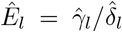 and *γ* = *Eδ*, then 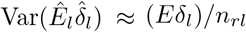. Now, by substituting *δ_l_* with 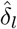, it follows that 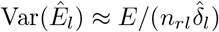. To minimize the variance of 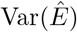, the inverse of 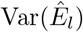 can be used as the weight to calculate the multi-locus estimate 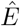. The unified weight *w_l_* is therefore equal to 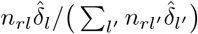, and 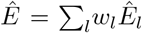. Because the loci are unlinked, we have 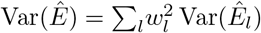, hence 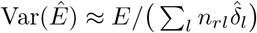.

The genotyping error rate *e* can now be estimated by the formula 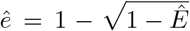. Moreover, because *e* ≈ *E*/2, the variance 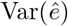 can be approximately expressed as 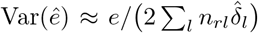. As described above, the inverse of 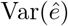 can be used to weight 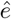 in multiple applications and datasets.

When the polyploid phenotypes are used, pair mismatches will be rare. Specifically, they are rare for the first category, because the single-locus exclusion rate is low (e.g. 0.01 for the hexaploid phenotypes at a hexa-allelic locus). Therefore, it is inaccurate to estimate *e* by pair mismatches. Relative to the first category, the single-locus exclusion rate for the second or the third categories is high (e.g. 0.27 for the hexaploid phenotypes at a hexa-allelic locus). Hence, we can use trio mismatches to reliably estimate the genotyping error rates for the second and the third categories, and the details are described in Appendix F.

### Estimation of sample rate

For an individual offspring, the probability that one of its true parents is sampled is defined as the *sample rate*, denoted by *p_s_*. The probability that an alleged parent (or a pair of alleged parents) of an offspring is assigned at a confidence level is called the *assignment rate*, denoted by *a*. Specifically, we denote *a_c_* for the assignment rate when the true parent(s) is sampled, and *a_u_* for the assignment rate when the true parent(s) is not sampled. Therefore, *a* is a weighted average of *a_c_* and *a_u_*.

We now develop a simple but robust estimator to estimate the sample rate from the assignment rate and begin our discussion with how to estimate the sample rate by using one application. For convenience, we will replace ‘the father’ with ‘one parent’ and ‘the mother’ with ‘the other parent’ in the first and the second categories in a parentage analysis.

For the first and the second categories, we have *a* = *p_s_a_c_* + (1 – *p_s_*)*a_u_*, so *p_s_* can be estimated by

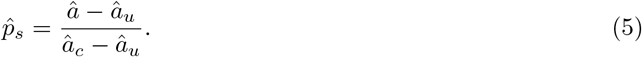

For the third category, if the sexes are known, then 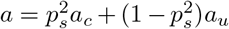, so *p_s_* can be estimated by

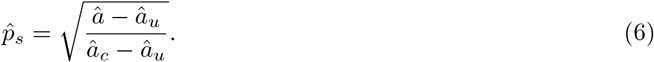

If the sexes are unknown, then *a* = *p_c_a_c_* + (1 – *p_c_*)*a_u_*, where *p_c_* is the probability that the true parents are sampled, which can be expressed as 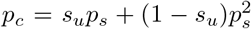, in which *s_u_* is the proportion of selfed offspring in this application. Hence 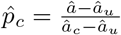, and the sample rate *p_s_* can be estimated by

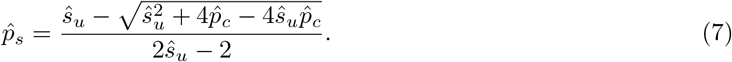

The value of 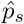 may be less than zero or greater than one. If this happens, we will truncate the value into the acceptable range [0, 1]. We will also set multiple confidence levels to estimate the selfing rate *s_u_* for increased accuracy. For the situations of multiple applications and multiple confidence levels, the estimation of the sampling rate is shown in Appendix G, along with the estimation of *s_u_*.

## Data Availability

polygene is written in C++ and C#, whose executables (Windows, Ubuntu and Mac OS X), source code and user manual are available on GitHub (http://github.com/huangkang1987/polygene).

The simulation functions are ‘`private void SIM_PARENT1()`’ to ‘`private void SIM_PARENT3()`’ in ‘`Forml.cs`’. The simulation parameters, output files, description of I/O format, figure plotting script and empirical dataset are available on the website of this journal.

## Evaluation

In this study, we use a computer simulation to create the genotypic and phenotypic datasets with disomic, tetrasomic or hexasomic inheritance, and then perform our parentage analysis by using these datasets. The performances of four methods under the same conditions are compared by four typical applications, where one method is the phenotype method, and the others are named the dominant method (Rodzen *et al*., 2004) (named after the pseudo-dominant data used in this method), the sibship method (Wang, 2016) (originating from the application ‘sibship reconstruction’) and the exclusion method (Zwart *et al*., 2016). The accuracies of these four methods under natural conditions are tested with an empirical microsatellite dataset for the highbush blueberry (Huber, 2016). In addition, the performances of the genotyping error rate estimation and the sample rate estimation are also evaluated using the simulated datasets.

Both the dominant and the sibship methods rely on first transforming the polyploid codominant phenotypic data into pseudo-dominant data. The same procedure as Kalinowski *et al*. (2007) is used for the dominant method, and the likelihood formulas under this method are listed in Appendix H, whose derivations are given by Gerber *et al*. (2000). Under the sibship method, a simulated-annealing algorithm is used to find the classification of optimal full-sib (or half-sib) families for the whole dataset by maximizing the likelihood, which is implemented in the software package colony (Wang and Scribner, 2014). Under the exclusion method, the effects of double-reduction and null alleles are incorporated, and the details of this method are described in Appendix I.

### Simulated data

In order to evaluate these methods, we create some theoretical monoecious populations, each consisting only of individuals with disomic to decasomic inheritance for the genotypic data or disomic to hexasomic inheritance for the phenotypic data. We assumed that the population under scrutiny is genotyped at *L* unlinked loci under the PES model (Huang *et al*., 2019). The number of loci *L* is set from three to 12 (genotypes) or three to 18 (phenotypes) at an interval of three. The distance (in centimorgans) between each of these loci and its corresponding centromere is drawn from the uniform distribution U(0, 100). The single chromatid recombination rate *r_s_* is obtained by Haldane’s mapping function. Each locus is located with six amplifiable alleles that have uniform initial frequencies, with the initial null allele frequency set as 0.1 for the phenotypic data. For the genotypic data, null alleles are not simulated because the dosage of alleles within each genotype is known.

Huang *et al*. (2019) derived the genotypic frequencies under each of the four double-reduction models listed in Table S1. However, the analytical solution of genotypic frequencies under inbreeding/selfing and double-reduction is still unknown. As an alternative, we give an approximated solution in Appendix C by using the inbreeding coefficient *F* as an intermediate variable with the assumption that any inbreeding is only caused by self-fertilization. With this approximation, we generate the genotypes of the founder generation by Equation (A4). In order to let the genotypic frequencies reach their equilibrium state and avoid severe genetic drift, 2000 individuals are generated for the founder generation, and the population is allowed to reproduce for ten generations, each generation consisting of 2000 individuals.

During reproduction, the parents of each offspring are either two distinct individuals randomly chosen from the previous generation at a probability of 1 – *s*, or the same individual (for self-fertilization) randomly chosen from the previous generation at a probability of *s*. The selfing rate *s* is set as three levels (0, 0.1 and 0.3). The following three procedures are designed to simulate meiosis: (i) the chromosomes are randomly paired and the alleles are exchanged between the pairing chromosomes at a probability of *r_s_*; (ii) the chromosomes are randomly segregated into two secondary oocytes; and (iii) the alleles within a chromosome are randomly segregated into two gametes. Fertilization is then simulated by the merging of two gametes.

Next, we reproduce two additional generations, each consisting of 100 individuals, to be used as the parents and offspring for the subsequent analyses. To simulate the missing parents, 90% of parents and all offspring are sampled. To simulate the genotyping errors, each genotype is swapped with the genotype of another individual at the same locus at a probability of 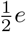 (where *e* is set as 0.01). To simulate negative amplification, each genotype is randomly set as ∅ at a probability of *β* (where *β* is set as 0.05). The phenotypes are obtained by removing both the null and the duplicated alleles within genotypes. Then the generated genotypic (or phenotypic) dataset is used to perform the parentage analysis. The allele frequency estimation is described in Appendix J.

For the first two categories in a parentage analysis, each is designated its own application (named Application (i) or (ii)). Application (iii) refers to a third category in which the alleged fathers and the alleged mothers are drawn from two different collections (representing that the sexes are known). Application (iv) also refers to the third category in which the alleged fathers and the alleged mothers are drawn from the same collection (representing that the sexes are unknown).

In Application (i), for each of the 100 offspring, 89 individuals from the parental generation are used as alleged fathers. Application (ii) is performed for the offspring with their mother sampled. In this application, for each offspring, the true mother is known, and 89 individuals from the parental generation are used as the alleged fathers. For Applications (i) and (ii), the alleged fathers will include the true father if sampled but will exclude the true mother (except the offspring is the product of self-fertilisation) to avoid interference. In Application (iii), for each offspring, 45 individuals (including the true father if sampled) from the parental generation are considered as the alleged fathers, with the remaining 45 individuals (including the true mother if sampled) as the alleged mothers. In Application (iv), for each offspring, all 90 individuals in the parental generation are considered as the alleged parents. We perform 100 replications for each of the three configurations: *v, L* and *s*, and calculate the average correct assignment rate for each configuration. Here, a *correct assignment* means that the true parents have been assigned and the value of Δ is higher than the corresponding threshold.

For the phenotype method, there are many models to estimate the allele frequencies and the related parameters, and the ideal way is to try each and then choose the optimal one with the smallest *Bayesian information criterion* (BIC) (as in Huang *et al*., 2020). However, it is time consuming to evaluate each of them in each simulation. As an alternative, we choose two models that work well in most situations: PES_0.25_ + *p_y_* + *β* + *s* for the phenotypic data and PES_0.25_ + *β* + *s* for the genotypic data. They denote the PES models with *r_s_* = 0.25 together with the considerations of null alleles (for phenotypes only), negative amplification and self-fertilization. Because the estimations of genotyping error rate *e* and sample rate *p_s_* depend on the number of assigned parents, the performance of a less efficient method will be reduced again due to the inaccurate estimations of *e* and *p_s_*. Since the aim of our simulation is to evaluate the performance of four methods, not the influence of the estimations of *e* and *p_s_*, the true values of *e* and *p_s_* are used as the *a priori* information. We perform 2000 Monte-Carlo simulations to obtain various critical values of the statistic Δ, and the correct assignment rates under three critical values (0, Δ_0.8_ and Δ_0.95_) are recorded.

For both the dominant and the sibship methods, the frequency *p*_dom_ of the dominant allele at a pseudo-dominant marker is estimated by 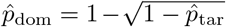, where 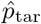 is the observed probability that a randomly sampled phenotype contains the target allele. For the dominant method, we implement the calculations of likelihood formulas listed in Appendix H in our simulation program. We also perform 2000 Monte-Carlo simulations to obtain the thresholds of Δ, and record the correct assignment rates under the same thresholds as above. For the sibship method, we write the pseudo-dominant phenotypes, the allele frequency estimates and other necessary parameters into a colony V2.0.6.5 input file. To avoid interference by the other cases, a unique input file for each case is generated. After calling `colony2p.exe` by a command-line mode, the results can be read from the output files. The probability of the identified parent(s) is used as a confidence level to compare with the phenotype and dominant methods. The exclusion method is implemented in our simulation program. In this method, the alleged parent (or parent-pair) with the fewest mismatches is assigned. If multiple alleged parents (or parent-pairs) have the same number of mismatches, none of them is assigned. For this method, any confidence level is unavailable.

For the four applications, each correct assignment rate as a function of *L* is denoted by a section of the overlapped bar charts, shown in Figure 1 for the genotypic data or in Figures 2, S1 and S2 for the phenotypic data.

**Figure 1.**
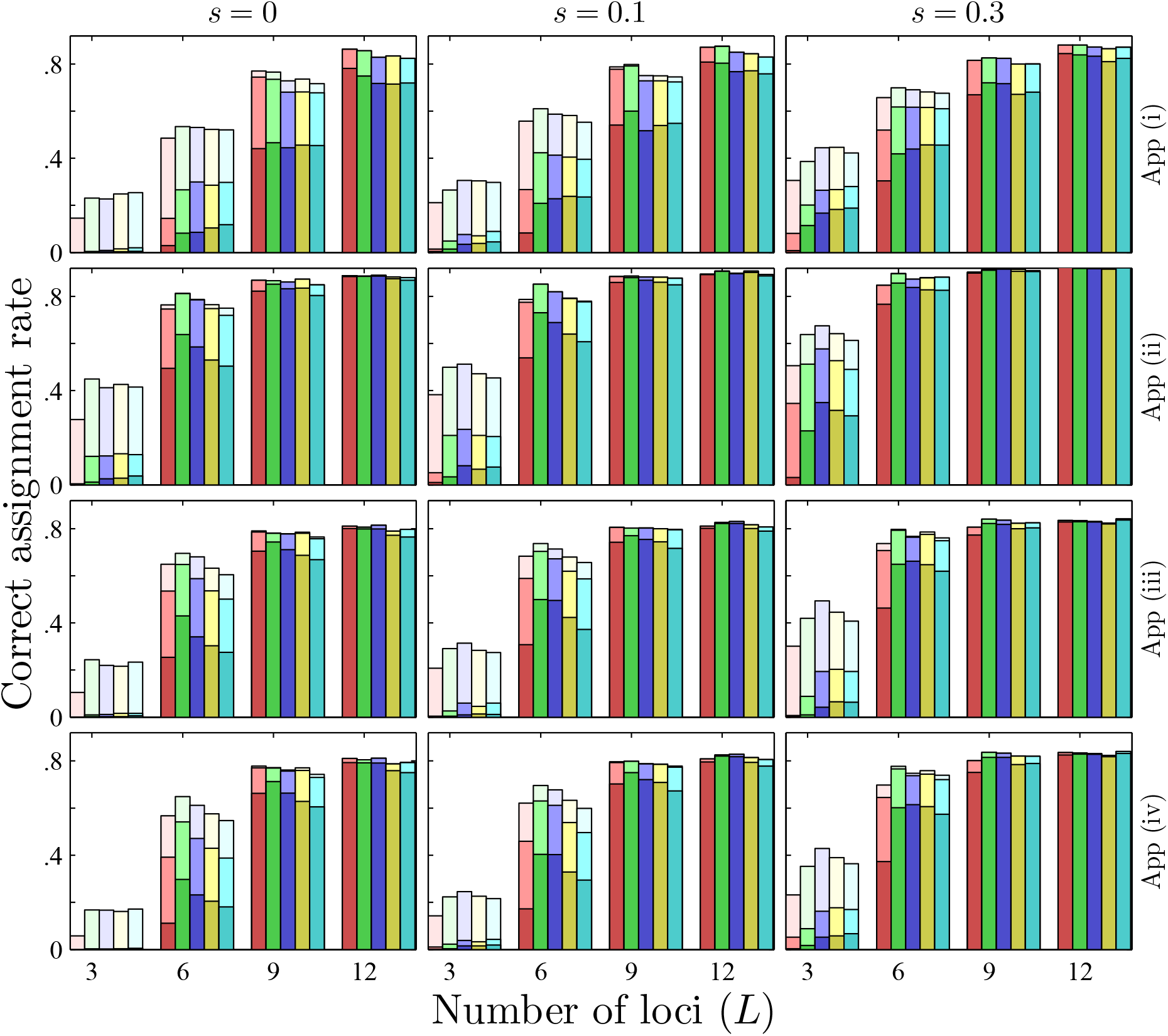
The correct assignment rate as a function of the number of loci *L* by using the genotypic data. Each row is designated an application and each column shows the simulation results for a different rate of selfing. Every correct assignment rate is denoted by a section of overlapping bar charts. The results of disomic to decasomic inheritances are shown by red, green, blue, yellow and azure bars, respectively. The bars with light, medium and bright colors denote in turn the correct assignment rates with the thresholds 0, Δ_0.80_ and Δ_0.95_.

**Figure 2.**
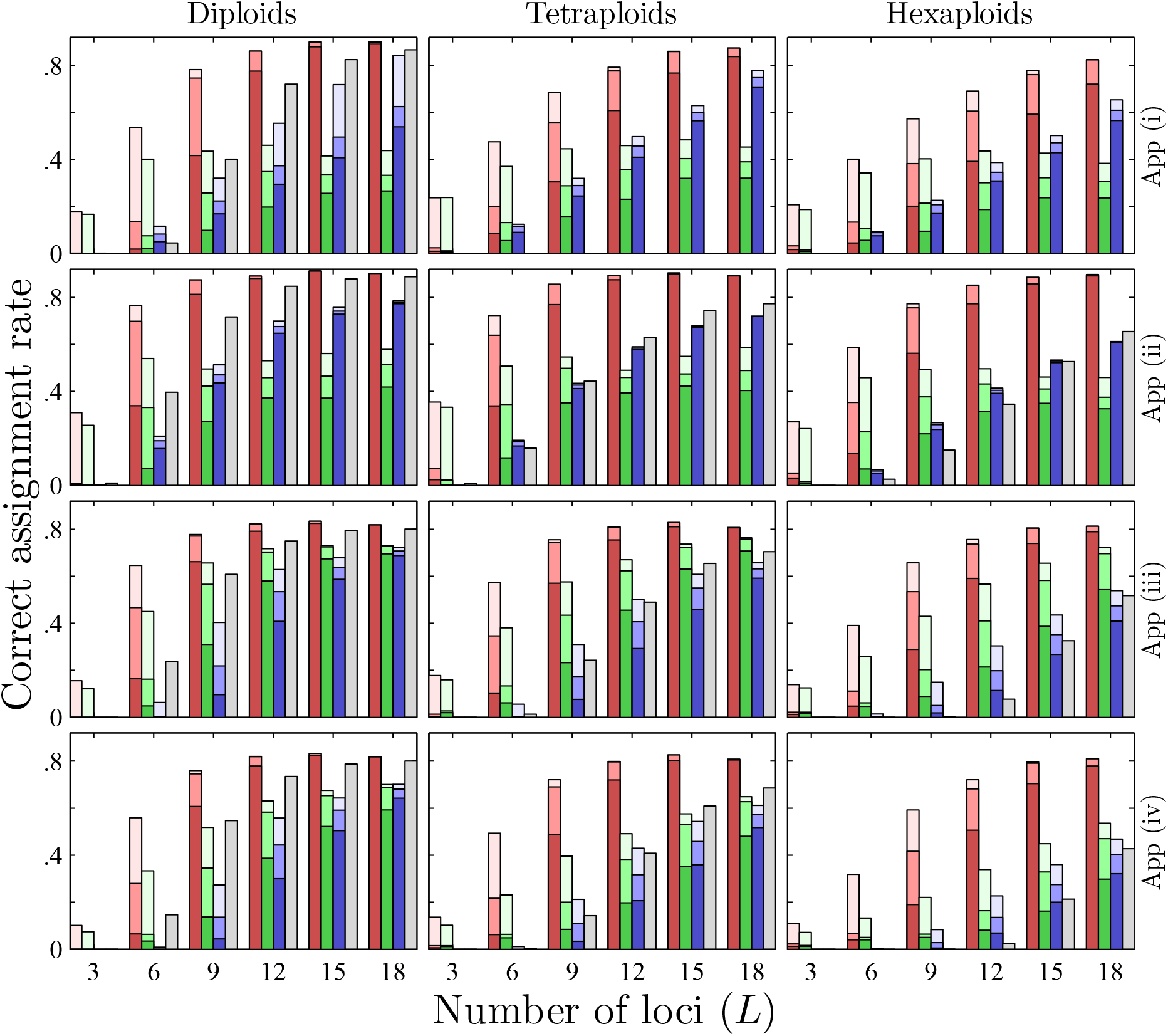
The correct assignment rates as a function of the number of loci *L* by using the phenotypic data at a selfing rate of 0.1. Each row is designated an application and each column shows the simulation results for a different ploidy level. The results for the phenotype, dominant, sibship and exclusion methods are shown by the red, green, blue and gray bars, respectively. The bars with light, medium and bright colors denote in turn the correct assignment rates with the confidence levels 0, 80% and 95%.

For the genotypic data, it can be seen from Figure 1 that each correct assignment rate increases as the number of loci *L* also increases, whose values reach a steady state if *L* is large enough (e.g. *L* ⩾ 12 for Application (i) or *L* ⩾ 9 for the other applications). The correct assignment rate generally reduces as the ploidy level increases. Moreover, as the selfing rate increases the correct assignment rate also increases but the difference among different ploidy levels decreases.

For the phenotypic data, it can be seen from Figure 2 that the correct assignment rate reduces as the ploidy level increases. The phenotype method outperforms the other methods, whose correct assignment rate at *L* = 9 is roughly the same as those of the other methods at *L* = 18, indicating that the phenotype method can reduce the number of loci needed to achieve the same accuracy by 40% to 60%. This method is also less sensitive to changes in the ploidy level, but an additional 23% and 45% loci are still required to reach the same correct assignment rate in tetraploids and hexaploids, respectively.

Compared with the dominant method, the performance of the sibship method is improved in Applications (i) and (ii) at a high *L* (⩾ 15), but is inferior in the other scenarios. The performance of the exclusion methods is good in Applications (ii) to (iv) at a high *L* (⩾ 15) but is inapplicable in Application (i).

It can be seen from Figures S2 and S3 that, like the results of genotypic data, the correct assignment rate is increased under most situations if the selfing rate is increased from 0 to 0.3. The assignment rate is reduced in Applications (ii) to (iv) under both the sibship and the exclusion methods.

### Empirical data

We used a microsatellite dataset from the highbush blueberry (*Vaccinium corymbosum*) (Chapter 5, Huber, 2016) to test the same four methods. The highbush blueberry has tetrasomic inheritance with no evidence of fixed heterozygosity (that indicates disomic inheritance; Krebs and Hancock, 1989).

The blueberry samples were collected from Agriculture Agri-Food Canada blueberry plots in Abbotsford and Agassiz, BC., Canada (Huber, 2016). Five controlled crosses, each with 25 to 30 offspring, were collected, resulting in a collection of 150 individuals, 143 of which were offspring. All samples were successfully amplified at 15 microsatellite loci, with the number of alleles sampled ranging from three to ten (Mean ± SD is 5.60 ± 2.33).

Following the four applications for the simulated data, we designed four similar applications for these empirical data. Application (I) or (II) refers to identifying the father when the mother is either unknown or known. There are 286 cases for each application, and each case has either 60 alleged fathers (including the true father and 59 false fathers) for Application (I) or the known mother together with 60 alleged fathers (including the true father and 59 false fathers) for Application (II). Application (III) refers to identifying the father and the mother jointly in which the alleged fathers and the alleged mothers are drawn from two different collections. There are 143 cases for this application, each of which has 30 alleged fathers (including the true father and 29 false fathers) and 30 alleged mothers (including the true mother and 29 false mothers). Application (IV) refers to identifying the father and the mother jointly in which the alleged fathers and the alleged mothers are drawn from the same collection. There are also 143 cases for this application, each of which has 60 alleged parents of unknown sex (including two true parents and 58 false parents).

There are altogether seven parents in these five controlled crosses. To increase the difficulty of our analysis, we also add 120 false parents which are generated by randomly copying the phenotypes from the real individuals. We randomly sample five to 15 loci from the dataset. For each value of *L*, 100 datasets are generated, each including 150 true individuals and 120 false parents. These datasets will be used to perform our parentage analysis by using the same four methods as described in the previous section. The analytical procedures are also the same as in the previous section except that the number of Monte-Carlo simulations to obtain the thresholds of Δ is 10,000 instead of 2000. The correct assignment rate will be used to measure the accuracy of each model.

The parentage assignment results from using each of the four methods and applying the phenotypic dataset of Huber (2016) are shown in Figure 3. The results patterns are similar to those obtained from the simulated data. The phenotype method still outperforms the other three methods but to a lesser degree than when the simulated dataset was used, but the phenotype method can still achieve the same accuracy with only 75% of the loci needed for the other methods. The exclusion method is still inaccurate and cannot be applied to real data in Application (I), but its performance is relatively good for the other applications when *L* > 10. The dominant method performs worse than the other three methods for Application (IV), as does the sibship method for Application (I).

**Figure 3.**
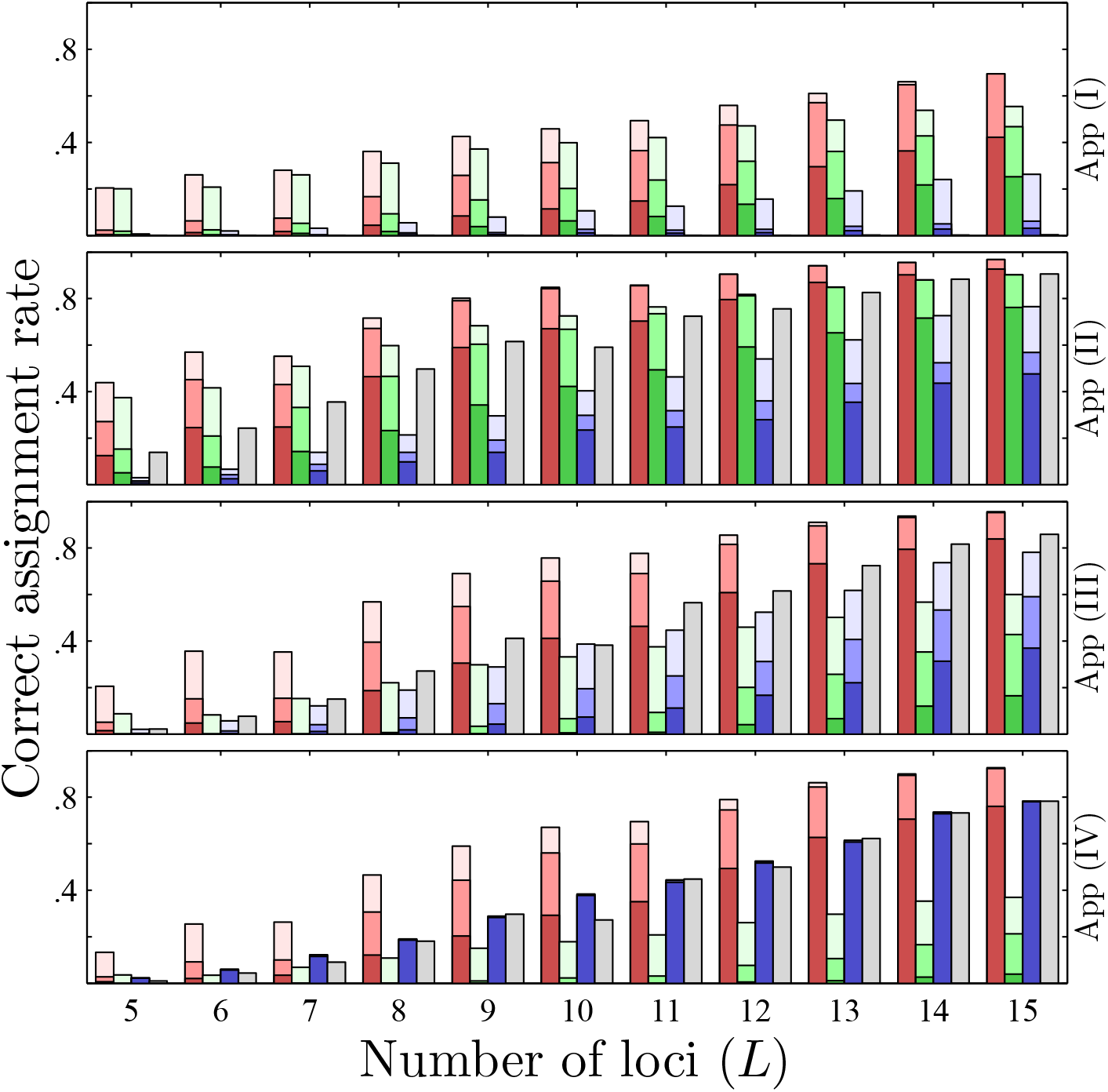
The correct assignment rates as a function of the number of loci *L* by using the phenotypic dataset of Huber (2016). Each row denotes an application. The methods, confidence levels and the definitions of bars together with their shading are as for Figure 2.

### Evaluation of genotyping error rate and sample rate

We use the simulated data to evaluate the performances of both estimators for the genotyping error rate and the sample rate. The same four applications are used as previously described, and are still referred to as Applications (i) to (iv). We estimate the genotyping error rate and the sample rate for each application. Two pairs of sampling and genotyping conditions, *poor* and *good*, are selected, which are *e* = 0.1 and *p_s_* = 0.5 for poor, or *e* = 0.02 and *p_s_* = 0.8 for good. The remaining parameters are almost the same as those in the section *Simulated data*, in which *s* = 0.1, *p_y_* = 0.1 and *L* is taken from six to 24 at an interval of three. We then perform 100 simulations for each configuration. The phenotype method is used to perform the parentage analysis with *a priori* genotyping error rate *e* = 0.01 and sample rate *p_s_* = 0.9. The allele frequencies are estimated under the PES_0.25_ + *p_y_* + *β* + *s* model. The performances of both estimators are evaluated by the RMSE.

For the estimation of the genotyping error rate, the identified pairs (or trios) with a confidence level of 99% are considered as the reference pairs (or trios), with *δ* estimated by randomly sampling 10,000 pairs (or trios). In Application (i), 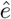 is estimated from the pair mismatch, whilst for the remaining applications 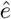 is estimated from the pair or the trio mismatches.

For the estimation of sample rate, we use the weighted average of 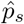 across three confidence levels (80%, 95% and 99%) for each application. Because 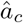 and 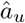 are obtained from the simulation, they may be influenced by any inaccurate simulation parameters, such as the sample rate, the selfing rate and the genotyping error rate. To improve the accuracy of these simulation parameters, we perform two rounds of analyses. The estimated sample rate and genotyping error rate in the first round are used as the *a priori* values in the second round. The results of the second round are used to evaluate the performance.

The results under both poor and good conditions are shown in Figures 4 and S4, respectively. For the estimation of the genotyping error rate, it can be seen that the results are good due to the RMSE being reduced to a low level. For example, the RMSE at *L* = 24 is able to reach 0.02 in poor conditions or 0.005 in good conditions. The RMSE for Application (i) performs worse than for the other applications, and increases greatly as the ploidy level also increases. This is because only the pair mismatch can be used for this application, and the single-locus exclusion rate for the first category is small. The RMSE for Application (ii) preforms better than for the other applications, because both the pair and the trio mismatches are used for this application, and the single-locus exclusion rate for the second category is usually higher than the other applications. The RMSE curves of Applications (iii) and (iv) are similar.

**Figure 4.**
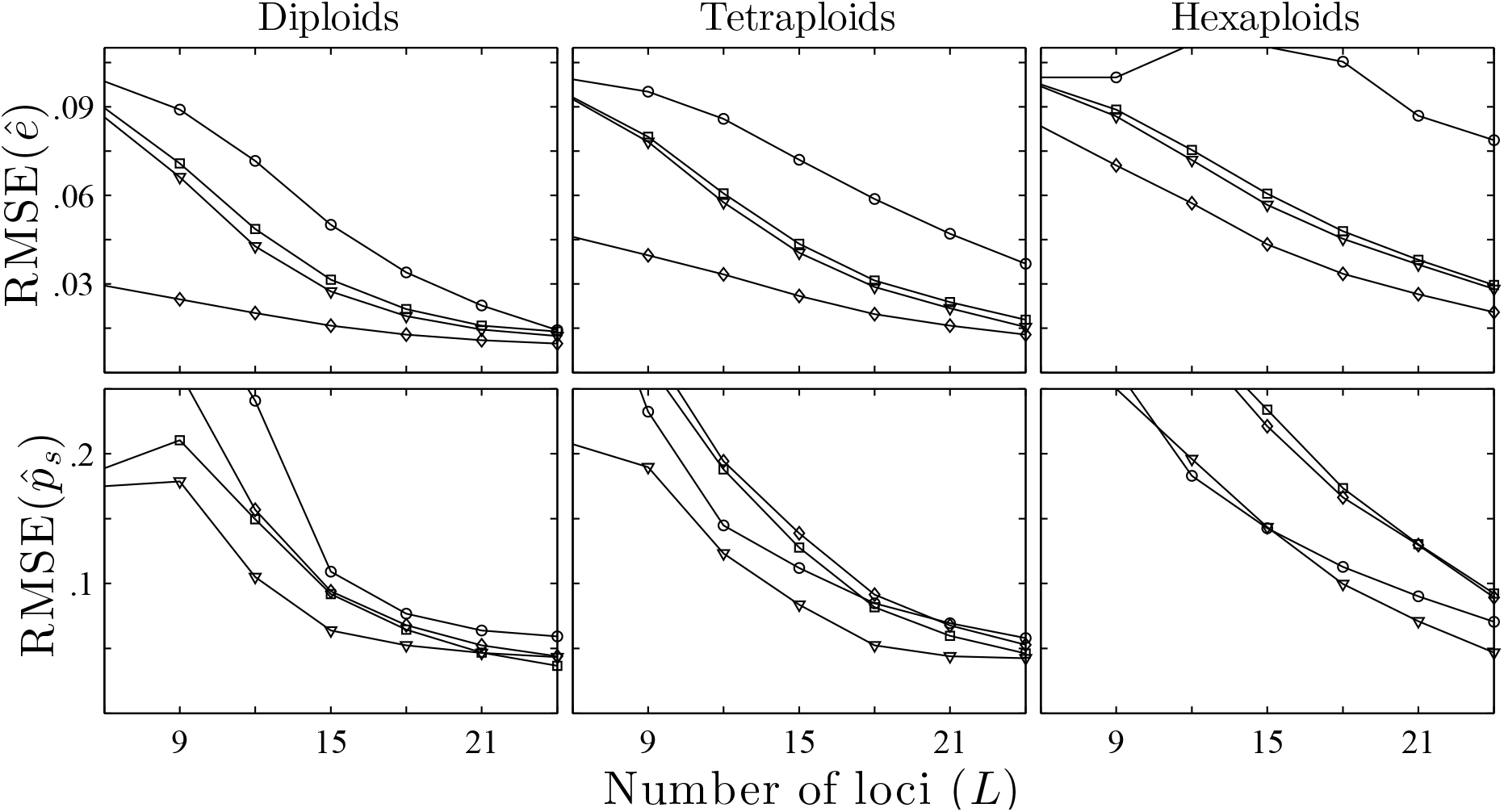
The RMSE of the estimated genotyping error rate 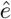 or the estimated sample rate 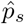 as a function of the number of loci *L* at *e* = 0.1 and *p_s_* = 0.5. Each column shows the results for a different ploidy level. The curves with circular, rhombic, triangular and squared markers denote the results for Applications (i), (ii), (iii) and (iv), respectively.

For the estimation of the sample rate, Figures 4 and S4 show that the results are inferior to those for the estimation of the genotyping error rate. For example, the RMSE at *L* = 24 is only able to reach 0.05 in poor conditions or 0.02 in good conditions. Unlike the estimation of the genotyping error rate, the results for Application (i) are not obviously inferior to those for the other applications. This is because the assignment rate rather than the reference pairs is used to estimate the sample rate, causing the results influenced less by the low single-locus exclusion rate.

The results for Application (ii) are poorer than those for estimating the genotyping error rate because fewer cases (≈ 50 cases) are used (about half of the true mothers are not sampled). If Applications (i) and (ii) use the same number of cases, then the performance of Application (ii) would be better than Application (i). Because Application (ii) also uses the mother’s data, which can better distinguish the true and the false fathers, the difference between *a_c_* and *a_u_* in Application (ii) is larger than that in Application (i) under the same conditions (e.g. Figure S3).

The results for Application (iii) are usually better than those for the other applications. This is because Application (iii) does not need to estimate the selfing rate and has a larger sample size (100 cases). However, the selfing rate has to be estimated for Application (iv), and thus the results are less accurate than for Application (iii).

## Discussion

### Inheritance model

Meiosis in polyploids is complex. Disomic and polysomic inheritances are two extremes, and many autopolyploid taxa represent the intermediate stages (Butruille and Boiteux, 2000). Allopolyploids (such as the segmental allopolyploids) can also display intermediate inheritance at some loci (Stift *et al*., 2008). In addition, some autopolyploid species can also form bivalent, univalent and other types of valents during meiosis (Lloyd and Bomblies, 2016). The formation of different types of valents may influence the sterility of the gametes or the seeds (Solís Neffa and Fernández, 2000)

For the autopolyploids with pure disomic inheritance, we can adopt the RCS model to simulate disomic inheritance. This is because the genotypic frequencies, gamete frequencies and transitional probabilities in the RCS model are the same as those for disomic inheritance. These probabilities are of interest for parentage analysis. The difference between the RCS model and disomic inheritance is that 100% multivalent formation is assumed in the former, whilst 100% bivalent formation is assumed in the latter. For the allopolyploids with pure disomic inheritance, all diploid methods including those of parentage analysis can be used if the genotypes at different isoloci are identified.

For intermediate inheritance, e.g. 50% bivalent and 50% multivalent gamete formation, regardless of how complex the nature of meiosis, identical-by-double-reduction (IBDR) alleles will be present in the resulting fertile gametes (Huang *et al*., 2019). For this reason, a generalized model was proposed, which uses ⌊*v*/4⌋ double-reduction rates in the calculation of genotypic frequencies and is able to describe meiosis patterns including that for intermediate inheritance (Huang *et al*., 2019). However, this model is too complex because it has ⌊*v*/4⌋ more degrees of freedom than the RCS (PRCS or CES) model. It is difficult to accurately estimate each double-reduction rate and thus is unrealistic to apply to many actual conditions. Even if these double-reduction rates are estimated, this model will often be suboptimal to other models because of the requirement for more degrees-of-freedom to explain various trends in a data set resulting in a higher BIC.

To better approximate the natural patterns, a simplified version of the generalized model was developed, named the PES model, which accommodates the single chromatid recombination rate *r_s_* as an additional parameter to calculate the genotypic frequencies (Huang *et al*., 2019). Especially, this model is equivalent to either the RCS model if *r_s_* = 0, or the CES model if *r_s_* = 1. Our software provides three PES-related models, which are the PES_0.25_, the PES_0.5_ and the PES estimate *r_s_*. The former two models do not increase their degrees-of-freedom because they use a fixed value of *r_s_*. We suggest to evaluate candidate models by the BIC and chose the optimal model with the lowest BIC (as in Huang *et al*., 2020).

### Performance of parentage analysis

For the genotypic data, the results for polyploids are generally similar to those for diploids (Figure 1). The correct assignment rate tends to increase if the ploidy level ranges from two to four, whilst the assignment rate decreases with a ploidy level that ranges from four to ten. However, this trend is weakened as the selfing rate increases.

These phenomena have at least three not necessarily mutually exclusive explanations. (i) At a high polyploid level, a genotype has more allele copies and so contains more genetic information (Huang *et al*., 2014). This can improve the performance of parentage analysis and many other population genetics analyses (e.g. the estimation of allele frequencies, genetic diversity, *F*-statistics, and relatedness coefficients). (ii) At a high polyploid level, the false parents are more likely to share the same alleles with the offspring, which may reduce the correct assignment rate. For example, if the ploidy level is high, reaching 1000, the false parents will share the same alleles with the offspring at a hexa-allelic locus. This is similar to when biallelic loci are used in tetraploids or hexaploids, the details of which are discussed in the following section. (iii) Selfing is able to reduce the difference among ploidy levels and improve the performance of our parentage analysis. Each of these three explanations will also be reflected in the phenotype results and are described at the end of this section.

For the phenotypic data, the results for polyploids are generally inferior to those for diploids for each application and for each method (e.g. see Figure 2). The phenotype method performs best among all four methods, saving at least 25% more loci than the other methods (e.g. see Figures 2 and 3), whose performances are stable for all applications.

For the four applications, the results of the phenotype method for diploids (Figures 2, S2 and S3) are slightly inferior to those for the genotypic data (Figure 1). This is because null alleles are simulated for the phenotypic data. In the absence of null alleles, each phenotype is only determined by one genotype for diploids. Therefore, both results under such condition are identical (data not shown).

For the dominant (Rodzen *et al*., 2004) and sibship (Wang and Scribner, 2014) methods, the results are suboptimal to those of the phenotype method (e.g. see Figures 3 and S2). In both the dominant and sibship methods, the polyploid codominant phenotypic data are transformed into the pseudo-dominant data, and the diploid procedures for a parentage analysis are subsequently used to perform an analysis. During transformation, genetic information is lost (Wang and Scribner, 2014) and some noise is also introduced. For example, in the pseudo-dominant approach the pseudo-dominant loci are assumed to be unlinked. In fact, because there are at most *v* alleles in a phenotype, the presence of an allele in a phenotype will reduce the probability of observing the other alleles in this phenotype, and so these loci are negatively correlated rather than unlinked. In addition, for the pseudo-dominant approach, many factors that affect the parentage analysis are not considered, such as double-reduction, null alleles, negative amplification, and inbreeding/selfing.

The exclusion method (Zwart *et al*., 2016) performs well in Applications (ii) to (iv), and the results are better than those for both the dominant and the sibship methods but only if *L* is high (e.g. see Figures 3 and S3). However, the exclusion method cannot be used for Application (i) because the single-locus exclusion rate in the first category is too low (e.g. 0.01 for hexaploid phenotypes at a hexa-allelic locus). Therefore, hundreds of loci are needed in order to exclude the false parents. This feature also influences the estimation of the genotyping error rate, such that the RMSE for Application (i) is highest (Figure 4).

From our simulation results, self-fertilization improves the accuracy of a parentage analysis, and reduces the variation of accuracies among different ploidy levels (Figures 1, 2, S2 and S3). This is because the genotypes become more homozygous as the selfing rate increases. If the selfing rate is one, all genotypes will become homozygotes at an equilibrium state. In such a case, each individual can be regarded as a haploid, and the ploidy level will not affect the accuracy of a parentage analysis.

### Genotyping error rate and sample rate

Our estimator for the genotyping error rate *e* is asymptotically unbiased as the number of loci increases. The bias of 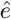 is from the estimation of *γ*. Because *γ* is estimated from any mismatches in the reference pairs or trios that are extracted from the identified parent(s), the confidence level of the true parents with few mismatches are successfully identified at a high probability. As a result, the value of 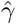 may be underestimated.

The estimation of the genotyping error rate does not use any simulation (*γ* is estimated from the reference pairs or trios, and *δ* is estimated from the distribution of the observed genotypes/phenotypes). This means that the estimator is not only robust but also insensitive to any errors in the simulation parameters (such as the allele frequency, negative amplification rate, selfing rate, sample rate, or the genotyping error rate). Any errors in these simulation parameters can only slightly affect the identified parents, which will not significantly affect the accuracy of 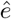. However, this estimator needs sufficient loci to identify the reference pairs or trios. For instance, if *e* = 0.1 and *p_s_* = 0.5, at least 15 loci are required in order to estimate the genotyping error rate for hexaploids in Application (i) (Figure 4).

Compared with the genotyping error rate, the estimation of the sample rate *p_s_* is less accurate and more sensitive to errors in the simulation parameters. There are at least three not necessarily mutually independent explanations for these patterns. (i) The estimate of the genotyping error rate is the weighted average of single-locus estimated values across all loci, where the actual sample size is 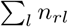. Whilst the sample rate is estimated only once for all loci, the actual sample size is the number of cases *n_c_* (see Appendix G). (ii) The sample rate estimator is biased in all categories in a parentage analysis because 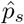 is truncated into the range [0, 1] and the operation of the square root is used in the third category. (iii) The simulation is used to obtain 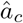 and 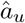 for the estimation of the sample rate, whilst the parameters used for simulation may be inaccurate (e.g. *a prior e* and *p_s_*). Any errors in 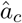 and 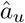 can be passed to 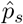, but such errors can be eliminated by increasing the number of loci. When the number of loci are sufficient, 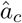 will be close to one, and 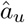 to zero. We suggest that users perform two rounds of estimation so as to reduce such errors as we have in the evaluation above.

### Polymorphism of loci

Because polyploids have more allele copies in a genotype, the false parents are more likely to share the same alleles with the offspring. Therefore, data resulting from the use of biallelic markers, e.g. *single nucleotide polymorphism* (SNPs), are unsuitable for performing a polyploid parentage analysis.

We will illustrate this by using the exclusion approach for the first category. For a given alleged parent, if its phenotype 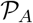 does not share any allele with its offspring phenotype 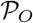, then it can be excluded as a true parent. If we assume that the double-reduction model is the RCS model, and that there are no interference factors (such as genotyping errors, self-fertilization, null alleles or negative amplification), then the exclusion rate Excl_1_ at a biallelic locus for the first category is

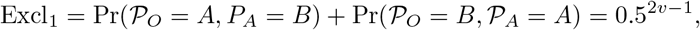

where *A* and *B* are the two alleles at this locus. The values of Excl_1_ from disomic to decasomic inheritances are in turn 0.125, 7.813 × 10^−3^, 4.883 × 10^−4^, 3.052 × 10^−5^ and 1.907 × 10^−6^. This sequence decreases exponentially, indicating that the false parents become less likely to be excluded as the ploidy level increases. Moreover, the number of loci required to achieve the combined exclusion rate 0.95 is ln(0.05)/ln(1 – Excl_1_), whose values from disomic to decasomic inheritances are in turn 22, 382, 6134, 98163 and 1570625.

Although next-generation sequencing (NGS) is able to segregate millions of SNPs, two reasons make it difficult to directly perform a parentage analysis with data obtained by using SNPs. First, the allele frequencies of most SNPs are not uniform, which reduces the exclusion rate. Second, adjacent SNPs are closely linked. This will reduce the accuracy of results because the genetic markers are assumed to be unlinked in all parentage analysis models.

Fortunately, haplotype assembly (Aguiar and Istrail, 2013), phased sequencing (Yang *et al*., 2011; Manching *et al*., 2017) and haplotype inference (Neigenfind *et al*., 2008) can all help to maintain the efficiency of NGS data, and can segregate multi-allelic loci by combining the closely linked variants so as to increase the single-locus polymorphism. Additionally, polyploid genotype calling can directly call back the genotypes but can currently only be applied to the biallelic variants (Carley *et al*., 2017; Weiß *et al*., 2018).

Multi-allelic markers can also be influenced by the same problem. We perform a simple simulation to describe the influence of the number of amplifiable alleles on the correct assignment rate, in which 20 loci with uniform amplifiable allele frequencies are used to perform our parentage analysis under the phenotype method (Figure 5). The correct assignment rate is much increased if the number of amplifiable alleles equates broadly to the ploidy level *v*, indicating that to achieve the optimal result, the number of amplifiable alleles should be greater than or equal to *v* (Figure 5). More loci are required if loci with relatively low levels of polymorphism are used. We suggest therefore to use highly polymorphic loci to perform parentage analysis.

**Figure 5.**
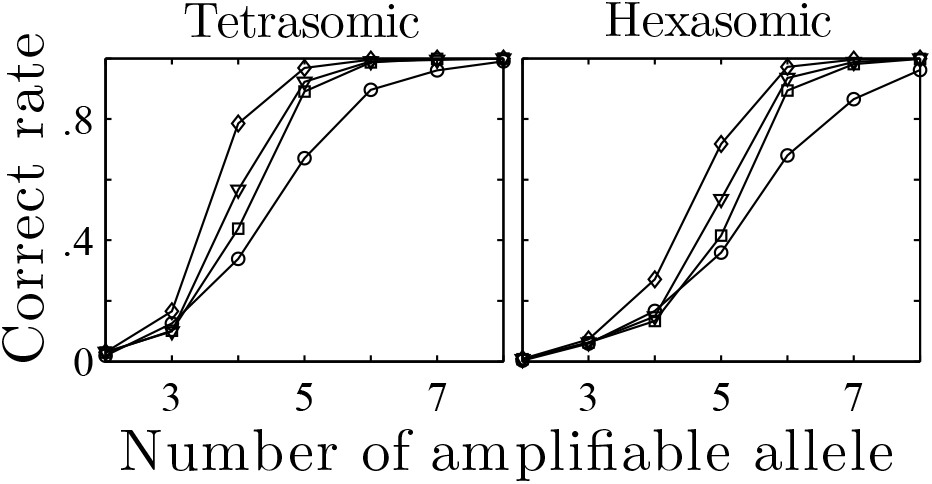
The correct assignment rates as a function of the number of amplifiable alleles under the phenotype method. Twenty loci with uniform allele frequencies of amplifiable alleles are used. The threshold and the selfing rate are set as Δ_0.95_ and 0.1, respectively. The remaining parameters and configurations are as for the simulated dataset. Each column shows the results for either tetrasomic or hexasomic inheritance. Each curve denotes the result for an application, whose definitions are as for Figure 4.

### Optimization and complexity

We use multi-threading, dynamic programming and genotype/phenotype indexing to optimize computational speed. The dynamic programming stores the likelihoods or LODs into a table so as to avoid repeated calculations. The genotype/phenotype indexing only records the hash values of genotypes /phenotypes for each individual, and the information of genotypes/phenotypes are saved in a hash table, that also includes the alleles, various frequencies (or prior/posterior probabilities), possible gametes and the number of occurrences.

All of these simulations took a total of three weeks to compute using a powerful workstation (Xeon E5 2699V4 36 cores). Computing efficiency will also be affected by the ploidy level *v* and the number of alleles *K* due to four main reasons: (i) the number of phenotypes increases as *v* and *K* increase, which reduces the efficiency of dynamic programming because more memory is required to store the likelihoods or LODs; (ii) the average number of genotypes determining a phenotype increases as *v* and *K* increase, which decelerates the calculation of likelihoods or LODs; (iii) the average number of gametes produced by a zygote increases as *v* and *K* increase, which decelerates the calculation of *T*(*G_O_* | *G_F_*) and *T*(*G_O_* | *G_F_*, *G_M_*) in Equation (A6); (iv) the number of terms in Equation (A7) increases as *v* and *K* increase, which decelerates the calculation of *T*(*g* | *G*) in Equation (A7). These four factors collectively and multiplicatively increase the complexity of the calculations. It is therefore not possible to perform an extensive simulation for highly polymorphic loci (e.g. *K* > 7) or for high ploidy levels (e.g. *v* = 8 or *v* = 10).

## Acknowledgements

We would like to thank the subject editor and two anonymous reviewers for their suggestions and comments. This study is funded by the Strategic Priority Research Program of the Chinese Academy of Sciences (XDB31020302), the National Natural Science Foundation of China (31730104, 31770411 and 31572278), the Young Elite Scientists Sponsorship Program by CAST (2017QNRC001), the National Key Programme of Research and Development, Ministry of Science and Technology (2016YFC0503200), and the Shaanxi Science and Technology Innovation Team (2019TD-012). Derek W. Dunn is supported by Shaanxi Province Talents 100 Fellowship.

## Author Contributions

KR and BGL designed the project, KH and KR constructed the model, GH provided the data, KH wrote the draft, GH and DD edited the manuscript.

## Supplementary materials

## Appendices

## A Double-reduction models

In the presence of double-reduction, a gamete will carry some *identical-by-double-reduction* (IBDR) alleles. For tetrasomic and hexasomic inheritances, there are only two and three allele copies within a gamete, respectively. Hence, there is at most one pair of IBDR alleles within a gamete. Therefore, we only need to use a single parameter to measure the degree of double-reduction.

For polysomic inheritance with a high ploidy level *v*, there may be more than one pair of IBDR alleles within a gamete. Therefore, it is necessary to add some additional parameters to measure the degree of double-reduction. Let *α_i_* be the probability that a gamete carries *i* pairs of IBDR alleles. Then 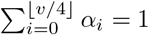, where ⌊*v*/4⌋ is the greatest integer not more than *v*/4. We call each *α_i_* a *double-reduction rate*.

Geneticists have developed several simplified models to simulate double-reduction. In the *random chromosome segregation* (RCS) model, the crossing over between the target locus and the corresponding centromere is ignored. Therefore, there cannot be any IBDR allele in a gamete, and the genotypic frequencies accord with the HWE (Figure S1(A), Muller, 1914).

The *pure random chromatid segregation* (PRCS) model accounts for such crossings over, and assumes that the chromatids behave independently in the meiotic anaphase, and are randomly segregated into some gametes (Figure S1(B), Haldane, 1930). When a pair of sister chromatids are segregated into the same gamete, the double-reduction occurs.

In the *complete equational segregation* (CES) model, the whole arms of two pairing chromatids are supposed to be exchanged between the pairing chromosomes (Figure S1(C), Mather, 1935). Subsequently, the chromosomes are randomly segregated into the secondary oocytes in Metaphase I. If the pairing chromosomes are segregated into the same secondary oocyte, the duplicated alleles may be further segregated into a single gamete.

The probability that an allele within a chromatid is exchanged with a pairing chromatid is called the *single chromatid recombination rate*, denoted by *r_s_*. In the CES model, the rate *r_s_* is assumed to be one. This is an ideal assumption. In fact, the maximum value of *r_s_* is 50% whenever the locus is located far from the centromere. Huang *et al*. (2019) presented a model by incorporating *r_s_* into CES, called the *partial equational segregation* (PES) model. Let *d* be the distance (in centimorgans) from the target locus to its corresponding centromere. According to the Haldane’s mapping function, the relational expression between *r_s_* and *d* is as follows:

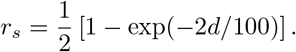

**Figure S1:**
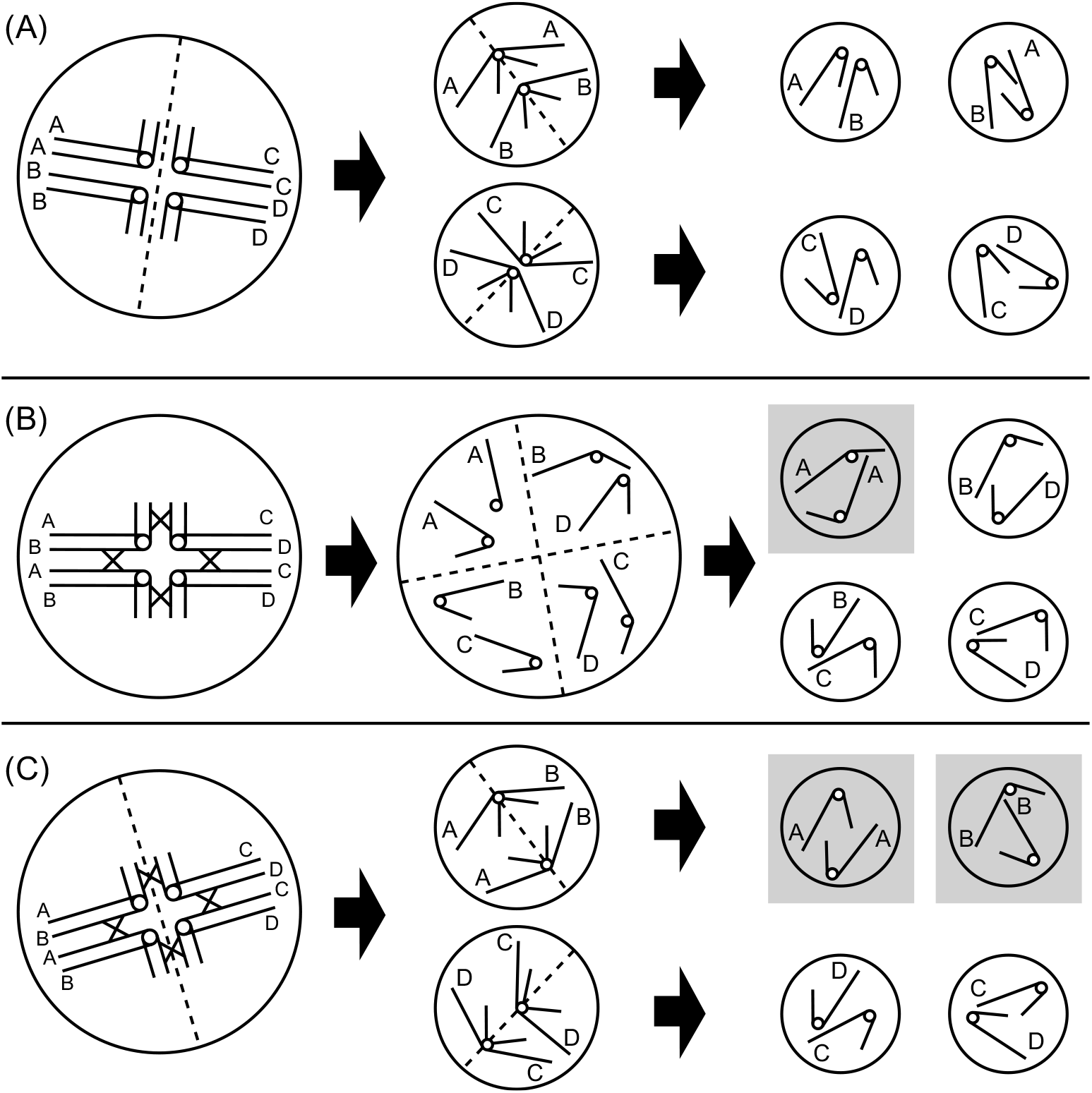
Diagram of double-reduction models under tetrasomic inheritance. The left column shows three primary oocytes, the middle column shows two secondary oocytes (in the rows marked (A) and (C)) or one tetrad (in the row marked (B)), and the right column shows three gametes. The gametes with a gray background carry IBDR alleles. We denote the cellular fissions by dashed lines, the arms of chromosomes by solid lines, and the centromeres by circles connecting solid lines. Each locus is located in a long arm of chromosomes and the identical-by-descent allele is denoted by the same letter as the corresponding locus. The row marked (A) is the sketch of RCS model. In this model, the crossing over between the target locus and its corresponding centromere is ignored (Muller, 1914). In the absence of crossing over, gametes may originate from any combination of homologous chromosomes, and two sister chromatids are never sorted into the same gamete (Parisod *et al*., 2010). The row marked (B) is the sketch of PRCS model. This model accounts for the crossing over between the target locus and its corresponding centromere, and assumes that the chromatids behave independently in the meiotic anaphase, and are randomly segregated into the gametes (Haldane, 1930). When a pair of sister chromatids are segregated into the same gamete, the double-reduction occurs. The probability that two chromatids within the same gamete are a pair of sister chromatids is 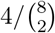, i.e. 1/7, where 4 is the number of pairs of sister chromatids, and 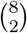 is the number of ways to sample two chromatids from eight chromatids. The row marked (C) is the sketch of CES model. In this model, the pairs of homologous chromosomes are exchanged with the chromatids via recombination (Mather, 1935). The whole arms of sister chromatids are exchanged into different chromosomes. The probability that two homologous chromosomes within a single secondary oocyte are previously paired at a locus in Prophase I is 1/3. In this case, the fragments of these sister chromatids will be segregated into a single gamete at the ratio of 1/2, so the double-reduction rate is 1/6 for tetrasomic inheritance.

In summary, different models are required to satisfy different conditions and their dimensions are also not the same. For example, there is an additional parameter *r_s_* (or *d*) in the PES model, and thus the number of degrees of freedom in PES is higher. It is noteworthy that all of the four models mentioned above can be incorporated into a generalized framework (i.e. the double-reduction rates are used as the parameters to express the phenotypic probabilities for some models). Comparing with the RCS, PRCS and CES models, the number of parameters for such generalized model increases by ⌊*v*/4⌋. The double-reduction rates in four models are shown in Table S1.

**Table S1:** The double-reduction rates in four models

| Model | Alpha | Ploidy level |  |  |  |  |
| --- | --- | --- | --- | --- | --- | --- |
|  |  | 4 | 6 | 8 | 10 | 12 |
| RCS | $\alpha_1$ | 0 | 0 | 0 | 0 | 0 |
| | $\alpha_2$ | | | 0 | 0 | 0 |
| | $\alpha_3$ | | | | | 0 |
| PRCS | $\alpha_1$ | 1/7 | 3/11 | 24/65 | 140/323 | 1440/3059 |
| | $\alpha_2$ | | | 1/65 | 15/323 | 270/3059 |
| | $\alpha_3$ | | | | | 5/3059 |
| CES | $\alpha_1$ | 1/6 | 3/10 | 27/70 | 55/126 | 285/616 |
| | $\alpha_2$ | | | 3/140 | 5/84 | 65/616 |
| | $\alpha_3$ | | | | | 5/1848 |
| PES | $\alpha_1$ | $r_s/6$ | $3r_s/10$ | $\frac{3}{70}r_s(10 - r_s)$ | $\frac{5}{126}r_s(14 - 3r_s)$ | $\frac{5}{616}r_s(84 - 28r_s + r_s^2)$ |
| | $\alpha_2$ | | | $\frac{3}{140}r_s^2$ | $\frac{5}{84}r_s^2$ | $\frac{5}{616}r_s^2(14 - r_s)$ |
| | $\alpha_3$ | | | | | $\frac{5}{1848}r_s^3$ |

## B Likelihoods for genotypic data

The likelihood formulas stated in this section are applicable to the genotypic data of both diploids and autopolyploids.

We will first give the likelihood formulas in the absence of self-fertilization, and these formulas are identical to those in Kalinowski *et al*. (2007). For the first category in a parentage analysis (i.e. identifying the father when the mother is unknown), the likelihoods can be expressed as

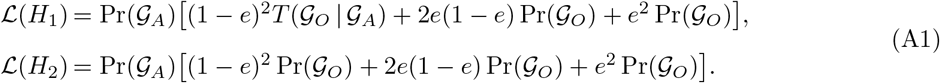

These two formulas are already listed in Equation (2), in which the second formula can be rewritten as 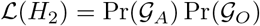 by merging similar terms.

For the second category (i.e. identifying the father when the mother is known), the likelihoods can be expressed as

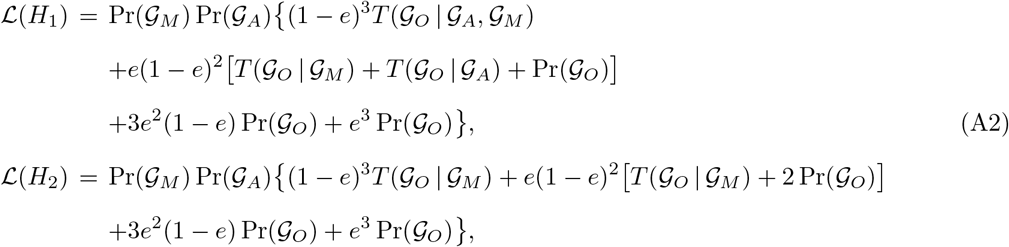

where 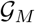 is the observed genotype of the true mother.

For the third category (i.e. identifying the father and the mother jointly), the likelihoods can be expressed as

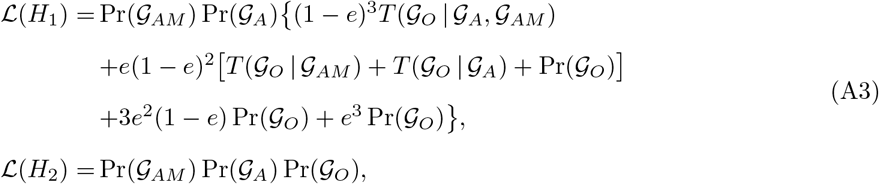

where 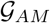 is the observed genotype of the alleged mother.

We will now give the likelihood formulas in the presence of self-fertilization. For the first category, the offspring is produced by selfing at a probability of *s* and by outcrossing at a probability of 1 − *s*. So, if we denote *T*_*s*1_ for 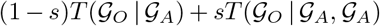, then the likelihood formulas can be obtained by replacing 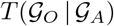 with *T*_*s*1_ in the first formula in Equation (A1), whose expressions are as follows:

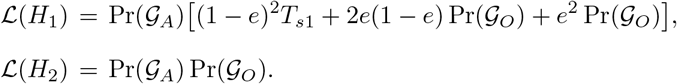

For the second category, if the alleged father is not the same individual as the true mother, selfing cannot occur in *H*_1_ but may occur in *H*_2_. Thus, if we denote *T*_*s*2_ for 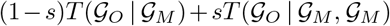, then the likelihood formulas can be obtained by replacing 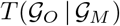 with *T*_*s*2_ in the second formula in Equation (A2), whose expressions are as follows:

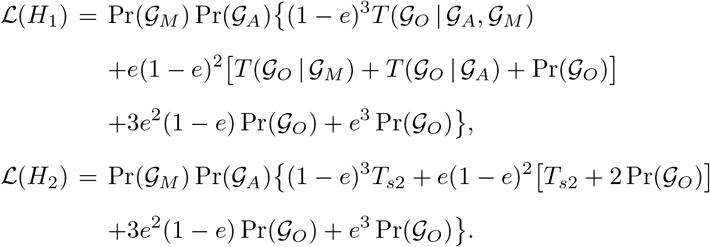

Moreover, if the alleged father is the same individual as the true mother, selfing must have occurred in *H*_1_ and could not have occurred in *H*_2_. Therefore, the likelihood formulas can be obtained by replacing 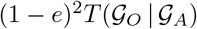 with 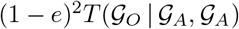 and 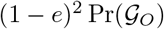 with 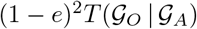 in Equation (A1), whose expressions are as follows:

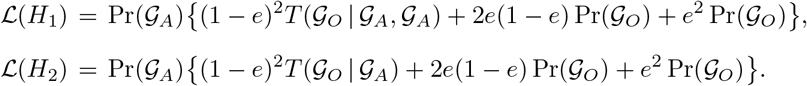

For the third category, if the alleged father is not the same individual as the alleged mother, selfing cannot happen in *H*_1_ but may happen in *H*_2_. In this situation, the likelihood formulas are the same as those in Equation (A3). Moreover, if the alleged father is the same individual as the alleged mother, selfing must have occurred in *H*_1_ but could not have occurred in *H*_2_. Therefore, the likelihood formulas can be obtained by replacing 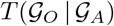 with 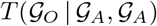 in the first formula in Equation (A1), whose expressions are as follows:

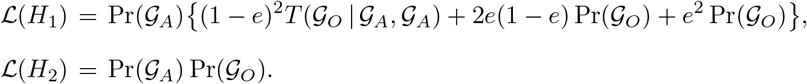

For the transitional probability 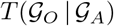 or 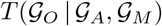 and so on in this section, it should be calculated by *T*(*G_O_* | *G_F_*) or *T*(*G_O_* | *G_F_, G_M_*) because these genotypes are assumed correctly genotyped in calculating these transitional probabilities, i.e. 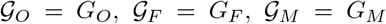. Similarly, for the genotypic frequency 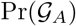 or 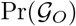 and so on in some formula listed in this section, it should be calculated by Pr(*G_A_*) or Pr(*G_O_*) because the genotyping errors does not change the distribution of genotypes, i.e. 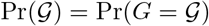.

For diploids without self-fertilization, the formulas of genotypic frequency and two transitional probabilities have been given in the section *Marshall et al.’s (1998) diploid model*.

For diploids with self-fertilization, the transitional probabilities do not change, but the genotypic frequency is related to the inbreeding coefficient *F*, denoted by Pr(*G* | **p**, *F*), which can be calculated by

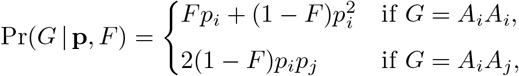

where *F* can be converted from the selfing rate *s* by the relational expression

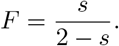

Above two formulas will be extended from disomic to polysomic inheritances in Appendix C.

For autopolyploids without self-fertilization, the genotypic frequency Pr(*G*) from tetrasomic to decasomic inheritances for each double-reduction model has been derived in Huang *et al*. (2019), and the transitional probabilities *T*(*G_O_* | *G_F_*) and *T*(*G_O_* | *G_F_, G_M_*) are given in Appendix D.

For autopolyploids with self-fertilization, the transitional probabilities do not change, but the exact genotypic frequency is unavailable. As an alternative, we give its approximate solution, whose derivation is given in Appendix C.

## C Genotypic and phenotypic frequencies

We have previously discussed the generalized genotypic frequencies from tetrasomic to decasomic inheritances under any double-reduction model (Huang *et al*., 2019). We will further incorporate self-fertilization into these genotypic frequencies.

In the presence of self-fertilization, if the ploidy level is high, the calculation of the genotypic frequencies from their analytical expressions is problematic (see Appendix K for details). As an alternative, we give an approximate solution by using the inbreeding coefficient *F* as an intermediate variable under the assumption that the inbreeding is only caused by both self-fertilization and double-reduction. The analytical expression of *F* at an equilibrium state under both double-reduction and selfing was derived in Huang *et al*. (2019), which is

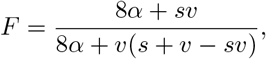

where *s* is the selfing rate, *v* is the ploidy level, and *α* is the expected number of pairs of IBDR alleles within a gamete. The value of *α* can be calculated by *α* Σ_*i*_ *iα_i_*, in which *α_i_* is a double-reduction rate, whose value is listed in Table S1.

Let’s now consider the genotypic frequencies incorporating both inbreeding and double-reduction. Let *p*_1_, *p*_2_, ⋯, *p_K_* be all allele frequencies in a population, and let *γ_k_* be (1/*F* − 1)*p_k_, k* = 1, 2, ⋯, *K*. Denote **p** = [*p*_1_, *p*_2_, ⋯, *p_K_*] and 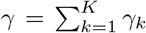. Assume that *q*_1_, *q*_2_, ⋯, *q_K_* are all allele frequencies of an individual, which are drawn from the Dirichlet distribution 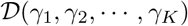 (Pritchard *et al*., 2000). Denote **q** = [*q*_1_, *q*_2_, ⋯, *q_K_*]. Then the probability density function of **q** is

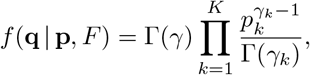

the expectation E(*q_k_*) is *p_k_*, and the variance Var(*q_k_*) is *Fp_k_*(1 − *p_k_*), *k* = 1, 2, ⋯, *K*. Moreover, for any *q_k_*, its standardized variance is exactly *F*. From this, we see that these conditions accord with those of the definition of Wright’s *F*-statistics. Hence the inbreeding coefficient *F* can be defined as the standardized variance of allele frequencies among individuals in the same population.

Because the correlation between alleles within the same individual relative to the population is explained by the divergence from **p** to **q**, the alleles within the same genotype are independent relative to **q**. Therefore, the frequency Pr(*G* | **q**) of a genotype *G* conditional on **q** is one of terms in the expansion of polynomial (*p*_1_ + *p*_2_ +⋯+ *p_K_*)^*v*^, i.e. the following term:

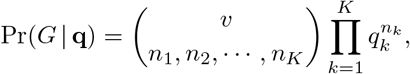

where *n_k_* is the number of copies of the *k*^th^ allele in *G, k* =1, 2, ⋯, *K*.

Next, the frequency Pr(*G* | **p**, *F*) of *G* conditional on **q** and *F* is the weighted average of all frequencies in the form of Pr(*G* | **q**), with *f*(**q** | **p**, *F*)d**q** as a weight, that is

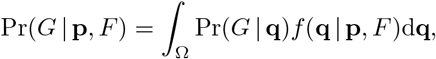

where the integral domain Ω can be expressed as

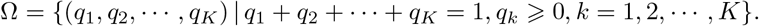

Such integral can be converted into the following repeated integral with the multiplicity *K* − 1:

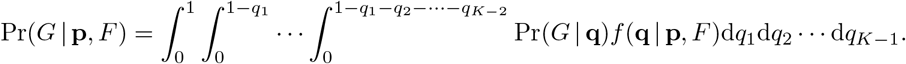

It can now be calculated from the expressions of Pr(*G* | **q**) and *f*(**q** | **p**, *F*) mentioned above that

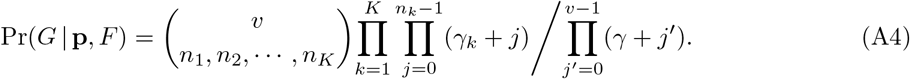

Equation (A4) is the approximate solution with *F* as an intermediate variable. Here, if self-fertilization is considered, the genotypic frequency 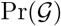 should be calculated by Equation (A4), otherwise, the formula of 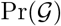 under each double-reduction model is given in Huang *et al*. (2019).

Based on the derivation above, we are now able to express the phenotypic frequencies whilst considering the presence of negative amplifications. If *β* is the negative amplification rate, the frequency 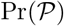 for each phenotype 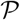 is the weighted average of 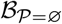 and 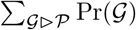 with *β* and 1 − *β* as their weights, i.e.

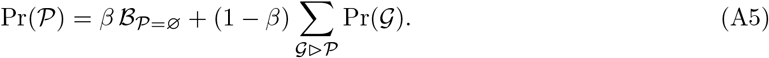

Besides, if the negative amplifications are not considered, it only needs to set *β* as zero in Equation (A5).

## D Transitional probabilities

In our model with a ploidy level greater than two, we establish two formulas of transitional probabilities *T*(*G_O_* | *G_F_*) and *T*(*G_O_* | *G_F_, G_M_*), whose expressions are as follows:

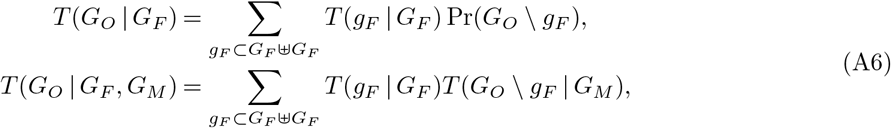

where the operations 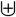 and \ are respectively the union and difference of multisets, *G_O_, G_F_* and *G_M_* are in turn the genotypes of the offspring, the father and the mother at a locus, *g_F_* and *G_O_* \ *g_F_* are the genotypes of the sperm and the egg that form the offspring, Pr(*G_O_* \ *g_F_*) is gamete frequency of the egg, and *T*(*g_F_* | *G_F_*) and *T*(*G_O_* \ *g_F_* | *G_M_*) are two transitional probabilities from a zygote to a gamete, which have been derived in Equation (A7).

It is noteworthy that there cannot be any double-reduction under the RCS model or the PES model with *r_s_* =0 (see Table S1), then the double-reduction should not be considered. In other words, the expression 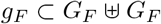 in Equation (A6) has to be replaced by *g_F_* ⊂ *G_F_* under these situations.

Huang *et al*. (2019) derived the generalized gamete frequency Pr(*g*) and zygote frequency Pr(*G*) (Huang *et al*., 2019). They also derived the generalized transitional probability *T*(*g* | *G*) from a zygote *G* to a gamete *g*, which can be used at any even ploidy level *v* and under any double-reduction model, whose expression is

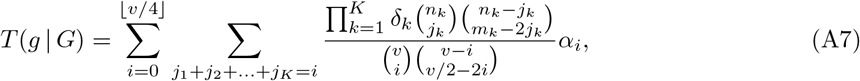

where *n_k_* (or *m_k_*) is the number of copies of the *k^th^* allele in *G* (or in *g*), *α_i_* is a double-reduction rate, and *δ_k_* is a binary variable, which is used to exclude the values outside the variation range *D* of *j_k_*, such that *δ_k_* = 1 if *j_k_* ∈ *D*, or *δ_k_* = 0 if *j_k_* ∉ *D*. The variation range *D* of *j_k_* can be expressed as

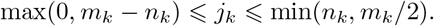

In fact, for the binomial coefficient 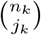, *n_k_* and *j_k_* should satisfy the condition 0 ⩽ *j_k_* ⩽ *n_k_*. Similarly, for 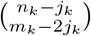, we have 0 ⩽ *m_k_* − 2*j_k_* ⩽ *n_k_* − *j_k_*, or equivalently *m_k_* − *n_k_* ⩽ *j_k_* ⩽ *m_k_*/2. Therefore, the expression of *D* holds.

## E Likelihoods under phenotype method

Under the phenotype method, if self-fertilization is not considered, the likelihoods for the first category in a parentage analysis can be expressed as

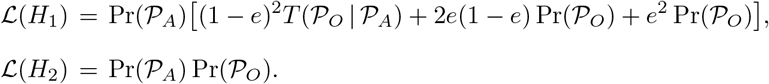

For the second category, the likelihoods can be expressed as

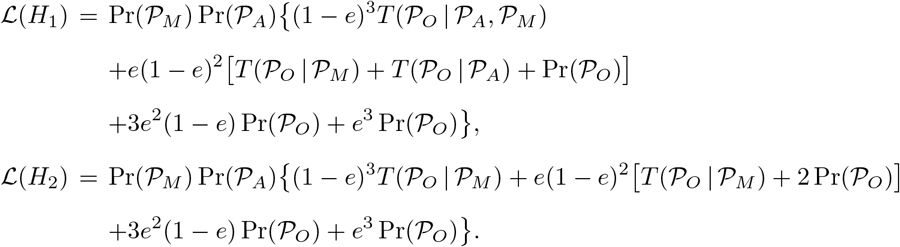

For the third category, they can be expressed as

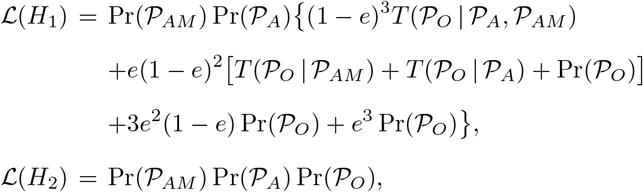

where 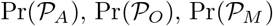 and 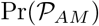 are calculated by Equation (A5), 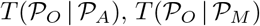 and 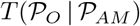 by Equation (3), and 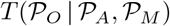 and 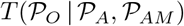 by Equation (4).

If self-fertilization is considered, like the situations of Appendix B, each pair of likelihood formulas can be obtained by modifying the existing formulas. For the first category, the likelihood formulas are

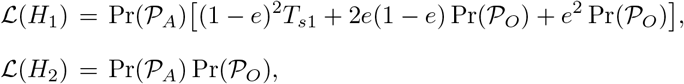

where 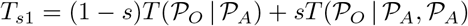. For the second category, if *A* ≢ *M*, then

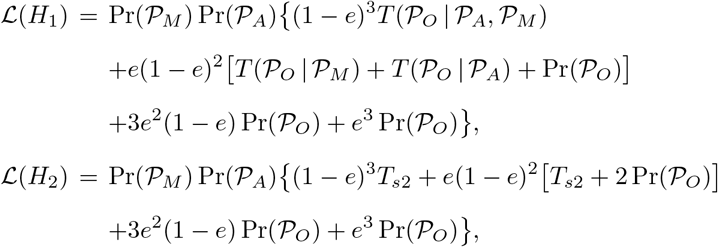

where 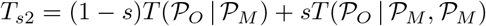; if *A* ≡ *M*, then

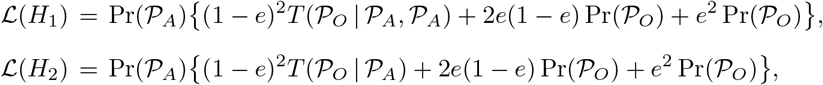

where 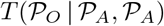 and 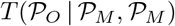 are calculated by Equation (4). For the third category, if *A* ≢ *AM*, then

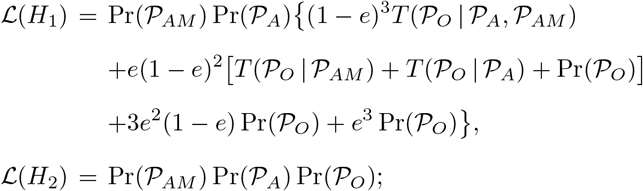

if *A* ≡ *AM*, then

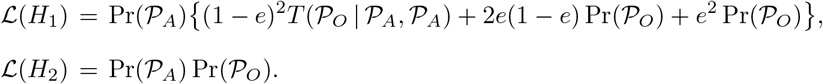

## F Estimation of genotyping error rate (continuous)

In this appendix, we will use the trio mismatches to describe how to estimate the genotyping error rate. The trio mismatch in a true parents-offspring trio may be caused by the genotyping errors in this offspring or in the parents. If the offspring or if both parents are erroneously genotyped, the probability of observing a trio mismatch is equal to the exclusion rate for the third category, denoted by *δ_o_*. If only one parent is erroneously genotyped, the probability of observing a trio mismatch is equal to the exclusion rate for the second category, denoted by *δ_p_*. Moreover, if each individual in a selfed trio is erroneously genotyped, the probability of observing a trio mismatch is denoted by *δ_s_*. Therefore, the probability *γ* of observing a trio mismatch in a true parents-offspring trio can be expressed as

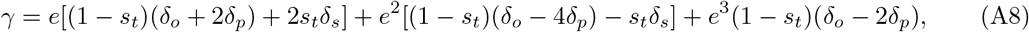

where *s_t_* is the frequency of selfing in the reference trios.

The values of *s_t_* and *γ* can be estimated from the reference trios identified from a single application or from multiple applications based on the same dataset, and *δ_o_* and *δ_s_* can be estimated from a similar Monte-Carlo algorithm mentioned above. The procedures are broadly as follows: randomly sample three (or two) individuals, considering them as a trio (or a selfed trio), and next calculate the probability that the genotypes/phenotypes at a locus of this trio (or this selfed trio) are mismatched, which is used as 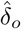 (or 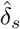) at this locus.

Under the assumption of random mating, the joint distribution of parental genotypes/phenotypes is the product of two observed genotypic/phenotypic frequencies, such that we can randomly sample two individuals and assume they are parents in the estimation of *δ_o_*. However, in the estimation of *δ_p_*, the joint distribution of parent-offspring genotypes/phenotypes cannot be estimated via this method. That is because the parent-offspring genotypes are correlated. As an alternative, we use the empirical distribution of genotypes/phenotypes of reference pairs to approximate the joint distribution of parent-offspring genotypes/phenotypes. More specifically, we randomly sample a matched pair (as a mother-offspring pair) from the reference pairs and an individual (as an alleged father) from all samples, considering them as a trio, and calculate the probability that the genotypes/phenotypes at a locus of this trio are mismatched, which is used as 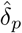 at this locus.

The single-locus estimate 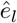 at the *l*^th^ locus can be obtained by solving Equation (A8), whose variance 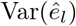 can be approximately expressed as 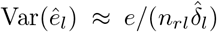. Moreover, the multi-locus estimate 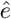 is the weighted average of single-locus estimates across all loci, that is 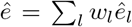, where 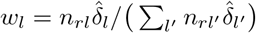. The variance 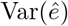 can be approximately expressed as 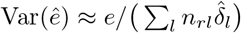.

## G Estimation of sample rate (continuous)

Assume that the assignment rates *a_c_* and *a_u_* as well as the selfing rate *s_u_* can be reliably estimated under an application and a confidence level, and that *n_c_* is the number of cases. Because the number of assigned cases *n_a_* obeys the binomial distribution B(*n_c_*; *a*), the assignment rate *a* can be estimated by 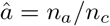. Therefore, the sample rate *p_s_* can be estimated by Equation (5), (6) or (7), and the variance 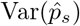 can be calculated by the formula 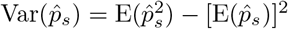.

However, it is unfortunate that the true value of *a* is unknown, then we cannot directly apply the binomial distribution B(*n_c_*; *a*) to perform various calculations. As an alternative, we select the uniform distribution U(0, 1) as the prior distribution obeyed by *a*, and then give the posterior distribution obeyed by *a* according to the Bayes formula, where the expected value E(*a*) for the posterior distribution is

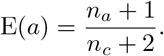

Now, we can perform various calculations so long as we let the value of *a* in B(*n_c_*; *a*) be equal to 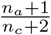.

In actual conditions, multiple applications and multiple confidence levels will be used jointly to increase the accuracy of sample rate estimation. For convenience, we denote 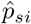 for the estimated value of *p_s_* under an application and a confidence level. According to the previous derivations, 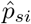 together with its variance can be calculated under the assumption that *a_c_, a_u_* and *s_u_* can be reliably estimated. Like the estimation of genotyping error rate, the estimate 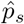 is the weighted average of the estimated values of *p_s_* under all selected applications and all selected confidence levels, symbolically 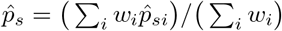, where 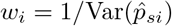.

Finally, let’s consider the estimation of selfing rate *s_u_* under multiple confidence levels. In actual conditions, the loci may be insufficient, causing that there are only few cases to assign the parent at a high confidence level (e.g. Δ > Δ_0.99_). Besides, the genotyping error rate may be high, causing that the false parent may be assigned at a low confidence level (e.g. Δ > 0) when the true parent is not sampled. To avoid these problems, we jointly use three confidence levels (80%, 95% and 99%) in polygene for each application.

The estimated value 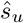 is the ratio of *n_s_* to *n_a_*, i.e. 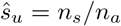 under an application and a confidence level, where *n_s_* is the number of selfing cases. If we select the three confidence levels 99%, 95% and 80%, then 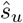 is the weighted average of the corresponding ratio values of *n_s_* to *n_a_*, that is

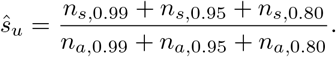

## H Pseudo-dominant approach

The pseudo-dominant approach was used in Rodzen *et al*. (2004) and Wang and Scribner (2014). In this approach, the codominant data are converted into the dominant data. More specifically, each visible allele is defined as a virtual dominant marker, whose observed phenotype is either present (denoted by {*A*}) if this allele is detected, or absent (denoted by ∅) if this allele is not detected. We denote 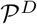 for the phenotype at a dominant marker. Moreover, the LOD scores are calculated by the diploid likelihood formulas listed below. These formulas are originally derived in Gerber *et al*. (2000) by using the transitional probability 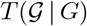 from a true genotype *G* to an observed genotype 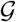 based on an alternative genotyping error model, where

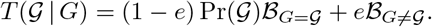

The above formula is different to that listed in Equation (1). Because the possible phenotypes at a dominant marker are {*A*} and ∅, the degree-of-freedom is only one. Therefore, the null allele frequency, the selfing rate and the negative amplification rate cannot be estimated. Besides, we will use the formulas and the model given in Rodzen *et al*. (2004) to evaluate the efficiency of this approach.

Next, the transitional probability from one phenotype or a pair of phenotypes to another phenotype at a dominant marker is described in Tables 1 and 2 in Gerber *et al*. (2000).

The phenotypic frequency at a dominant marker in diploids is

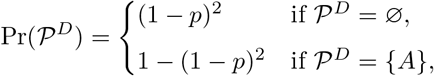

where *p* is the frequency of the dominant allele *A* at this dominant marker, and *p* is estimated from the observed phenotypic frequencies, whose estimated expression is 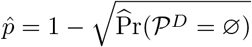.

Now, the likelihood formulas listed below can be used for the actual calculation by using these transitional probabilities and phenotypic frequencies: for the first category in a parentage analysis,

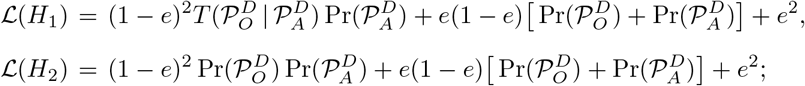

for the second category,

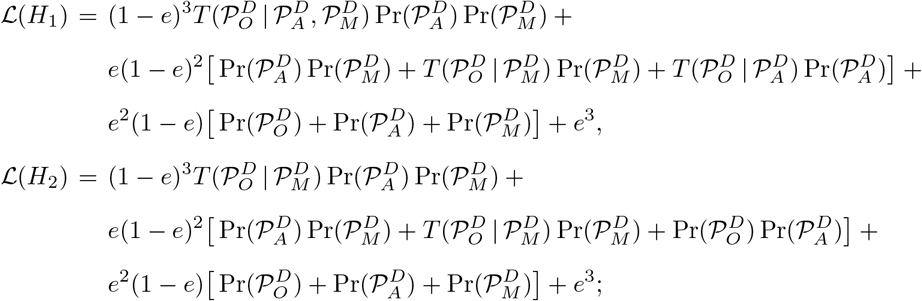

for the third category,

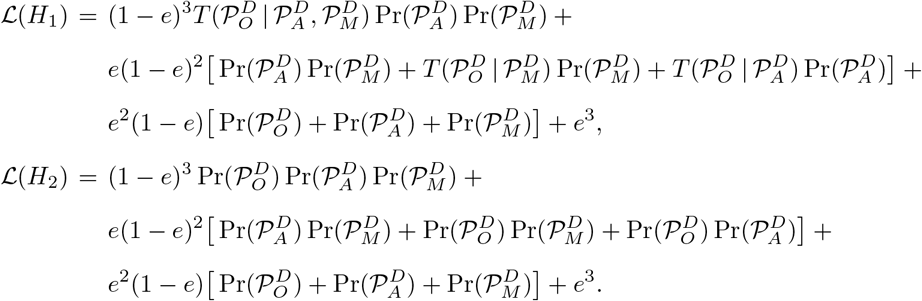

## I Exclusion approach

Although the exclusion approach is not as accurate as the likelihood approach, the number of mismatches can be used as a reference. Here, we extend the exclusion approach to polysomic inheritances, and this extended approach can be incorporated into our framework, such that the effects of double-reduction, null alleles, negative amplifications and self-fertilization can all be freely accommodated.

The logic of the exclusion approach is relatively simple: if the alleged parents are able to produce the offspring, they cannot be excluded. We will here give two extended definitions of matches by using the genotypic data.

Given an alleged parent-offspring pair, if there exists a gamete *g_A_* produced by the alleged parent at a locus, such that *g_A_* is a subset of the offspring genotype 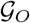 at this locus, then such a pair is termed *matched* at this locus. The condition in this definition can be described by symbols as follows: 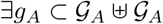, such that 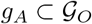; or equivalently, 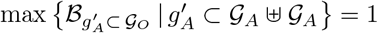, where 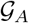 is the genotype of the alleged parent at this locus.

Given an alleged parents-offspring trio, if there exist two gametes *g_F_* and *g_M_* produced by the alleged father and the alleged mother at a locus, respectively, such that the fusion of *g_F_* and *g_M_* results in the offspring genotype 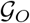 at this locus, then such a trio is termed *matched* at this locus. Similarly, the conditions in this definition can be described as follows: 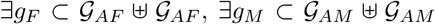, such that 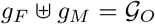; or equivalently,

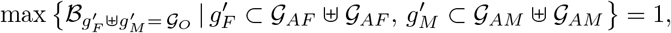

where 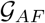 (or 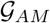) is the genotype of the alleged father (or the alleged mother) at this locus.

Finally, it is important to highlight that under the RCS model or the PES model with *r_s_* = 0, the expressions, used to describe the two definitions and involved in the double-reduction, should be revised, i.e. we must replace 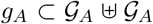 by 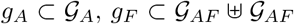 by 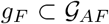 and 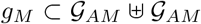 by 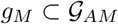.

## J Allele frequency estimation

We adopt an *expectation-maximization* (EM) algorithm (Dempster *et al*., 1977) to estimate the allele frequencies for phenotypic data. This algorithm follows the methods of Kalinowski and Taper (2006), which is an iterative algorithm used to maximize the genotypic likelihood. The *genotypic likelihood* at a locus is defined as the product of genotypic frequencies of all individuals at this locus, denoted by 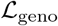, whose logarithmic expression is

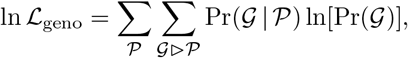

in which 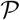 is taken from the phenotypes of all individuals at this locus, 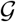 is taken from all genotypes determining 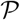 at the same locus, 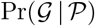 is the posterior probability of 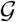 determining 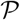, and 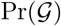 is the frequency of 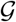.

The initial frequencies of amplifiable alleles are assumed to be equal to 1/*K*, where *K* is the number of alleles, including the null allele *A_y_*. The updated frequency 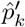 of the *k*^th^ allele *A_k_* is the weighted average of frequencies of *A_k_* in all genotypes at a locus, with the posterior probabilities of these genotypes as their weights, whose expression is

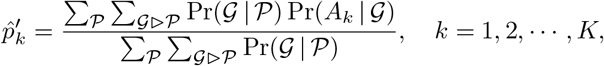

where 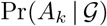 is the frequency of *A_k_* in 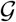.

Our algorithm also includes simultaneously the estimation of negative amplification rate *β*. Because the final estimated value of *β* is independent to the initial value, the initial value can be arbitrarily selected (e.g. 0.05). The updated negative amplification rate 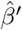 can be expressed as

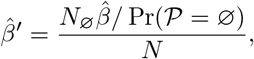

where *N*_∅_ is the number of negative phenotypes at this locus, *N* is the number of all individuals, 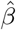 is the current negative amplification rate, and 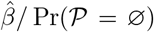 is the posterior probability that a negative phenotype is the result of negative amplification.

If 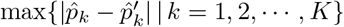 and 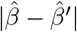 are less than a predefined threshold (e.g. 10^−5^) or if the iterative times reach 2000, the iteration is terminated, where 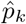 is the current frequency of *A_k_*.

Null alleles and negative amplifications can both be freely incorporated into our model. If the null alleles are not considered, the candidate genotypes extracted from a phenotype only need to be set as ‘not containing *A_y_*’. If the negative amplifications are not considered, the initial value of *β* only needs to be set as zero. If both factors are not considered, the negative phenotype cannot be explained, and so ∅ is discarded in the allele frequency estimation together with the subsequent analyses.

We also nest a downhill simplex algorithm (Nelder and Mead, 1965) outside the EM algorithm to estimate the selfing rate *s*. The estimated value *ŝ* is obtained by maximizing the phenotypic likelihood 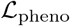, that is 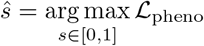, where 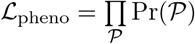.

## K Reasons for computational difficulty

In the absence of selfing, the generalized form of genotypic frequencies can be obtained by two methods (Huang *et al*., 2019). The first method is the *non-linear method*. In this method, we establish a non-linear equation set with the frequencies Pr(*G*_1_), Pr(*G*_2_), ⋯, Pr(*G_I_*), Pr(*g*_1_), ⋯, Pr(*g_J_*) as the unknowns and the frequencies *p*_1_, *p*_2_, ⋯, *p_K_* as the parameters, whose expression is as follows:

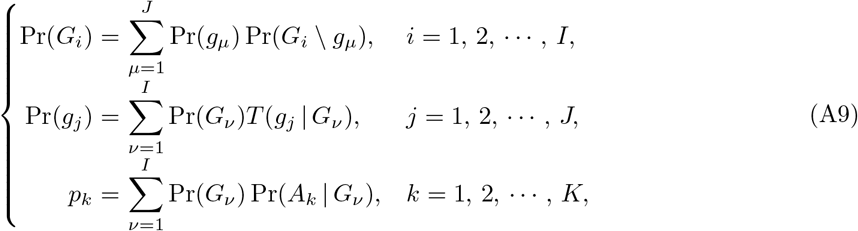

where 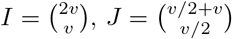, *K* = *v* + 1 (*I, J* and *K* are the numbers of zygotes, gametes and alleles at a locus, respectively), Pr(*G_i_* \ *g_μ_*) = Pr(*g* = *G_i_* \ *g_μ_*), *T*(*g_j_* | *G_ν_*) is the transitional probability from *G_ν_* to *g_j_*, and *p_k_* and Pr(*A_k_* | *G_ν_*) are the frequencies of *A_k_* in the population and in *G_ν_*, respectively. If the ploidy level *v* is equal to 4, 6, 8 or 10, the number of equations in Equation set (A9) is 90, 1015, 13374 or 187770, and the number of unknowns is 85, 1008, 12265 or 187759. We now see that these numbers will increase rapidly with an increase in ploidy level. Therefore, this will cause a computational difficulty for Equation set (A9) at a high ploidy level.

In order to overcome such a computational difficulty, we adopt another method, named the *linear method*, to obtain the zygote frequencies. For this method, briefly speaking, we will first use Equation set (A9) to calculate the gamete frequencies at a biallelic locus. Next, we split these alleles one by one at this locus until they are split into *v*/2 + 1 alleles so as to more expediently obtain the zygote frequencies at a multi-allelic locus. Finally, we use the former *I* equations in Equation set (A9), i.e.

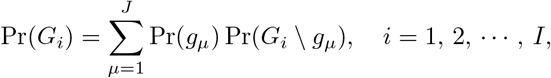

to calculate the zygote frequencies. This method can be described by a linear equation set **Ax** = **b**. Because there are no sufficient constraint conditions to obtain a unique solution for such linear equation set when *v* ⩾ 12, this method can only be applied from tetrasomic to decasomic inheritances (Huang *et al*., 2019).

In the presence of selfing, for the linear method, although the gamete frequencies can be solved for *v* < 12, the zygote frequencies cannot be easily calculated from the gamete frequency. That is because for any *i* ∈ *I*, the *i*^th^ equation in Equation set (A9) should be modified as

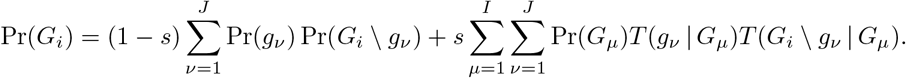

For the non-linear method, the calculation is more difficult when the ploidy level is high.

## L Supplementary figures

**Figure S2:**
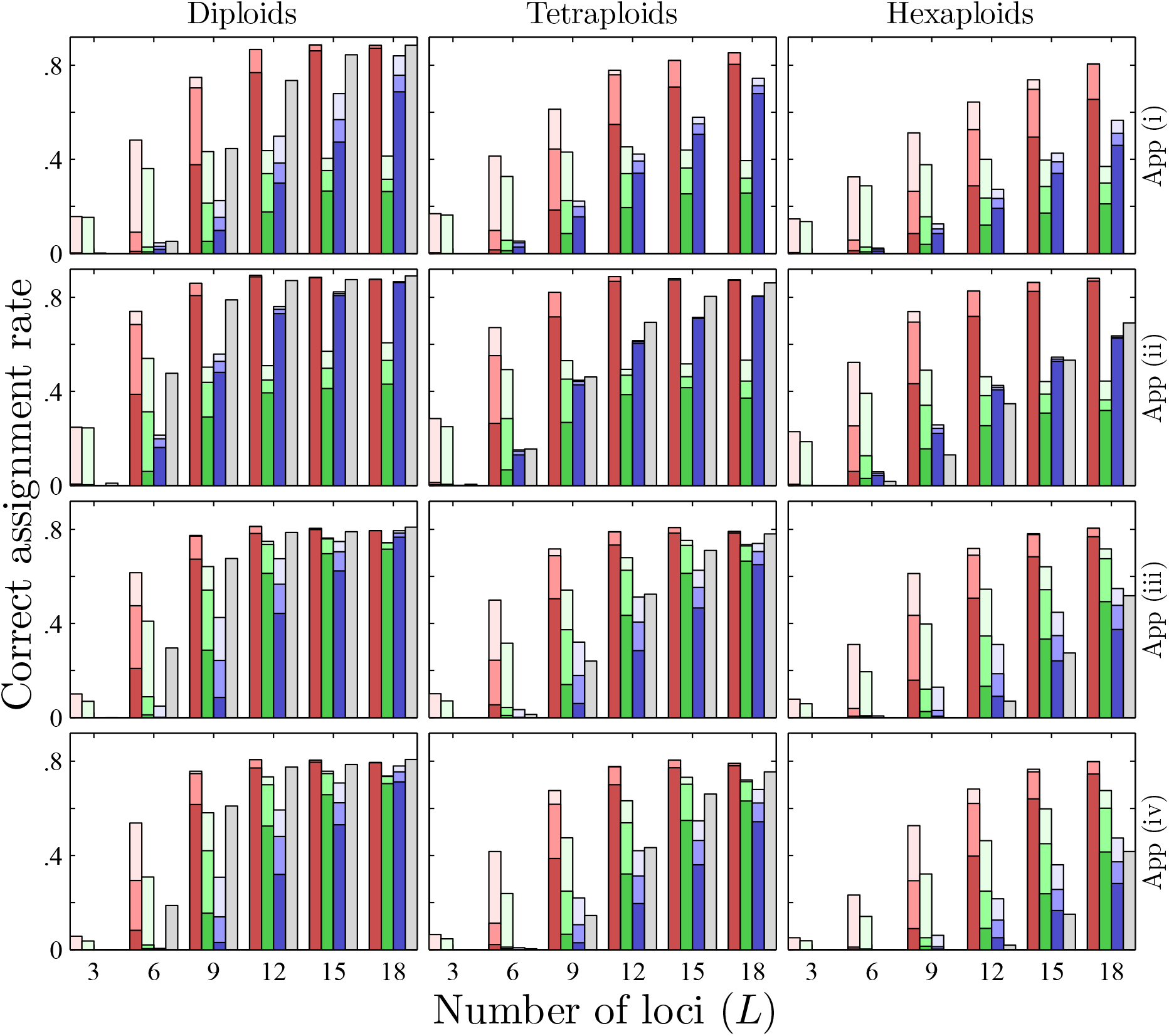
The correct assignment rate as a function of the number of loci *L* by using the phenotypic data at the selfing rate 0. The ploidy levels, applications, methods, confidence levels and the definitions of bars together with their shading are as for Figure 2.

**Figure S3:**
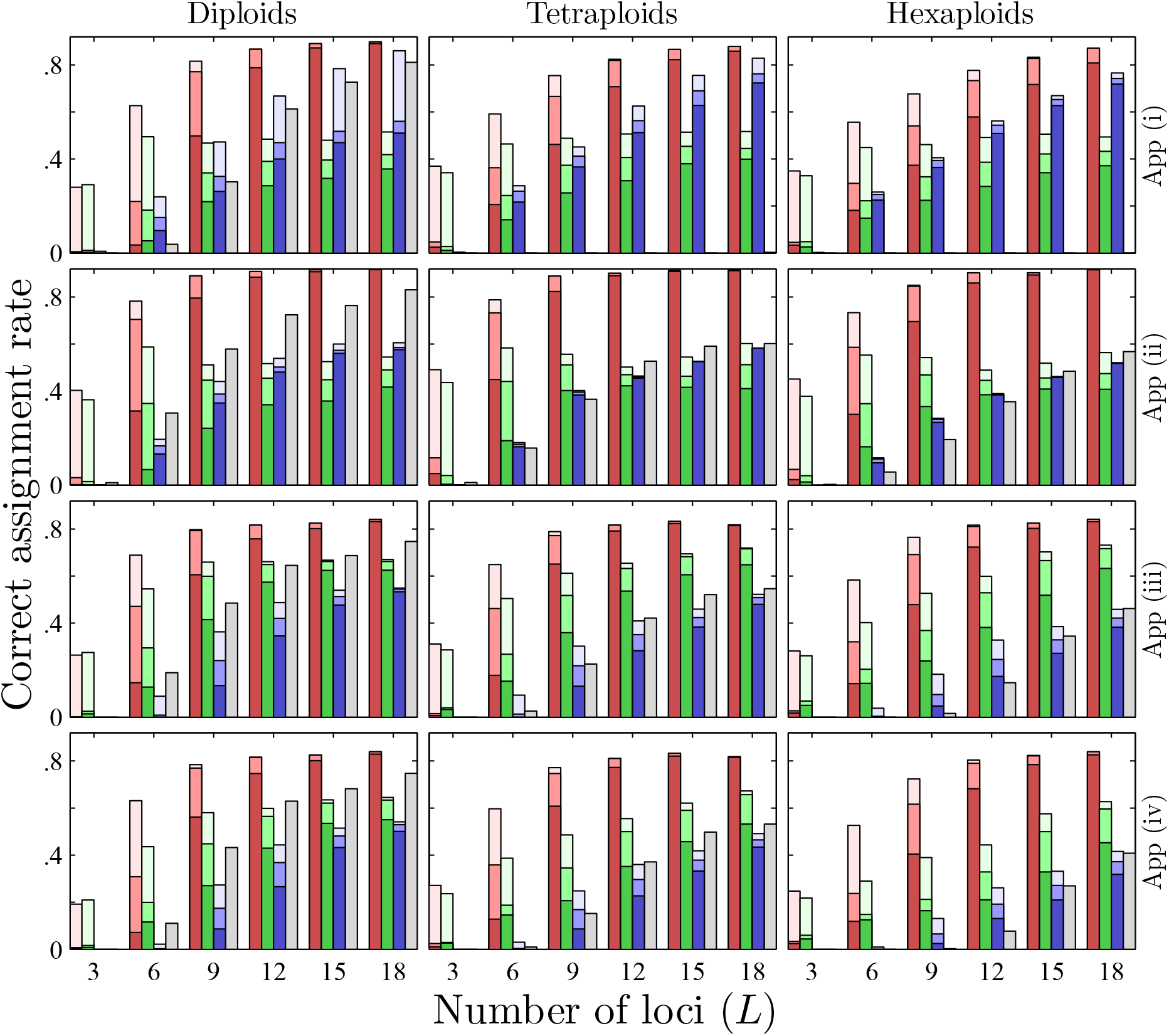
The correct assignment rate as a function of the number of loci *L* by using the phenotypic data at the selfing rate 0.3. The remaining are as for Figure 2.

**Figure S4:**
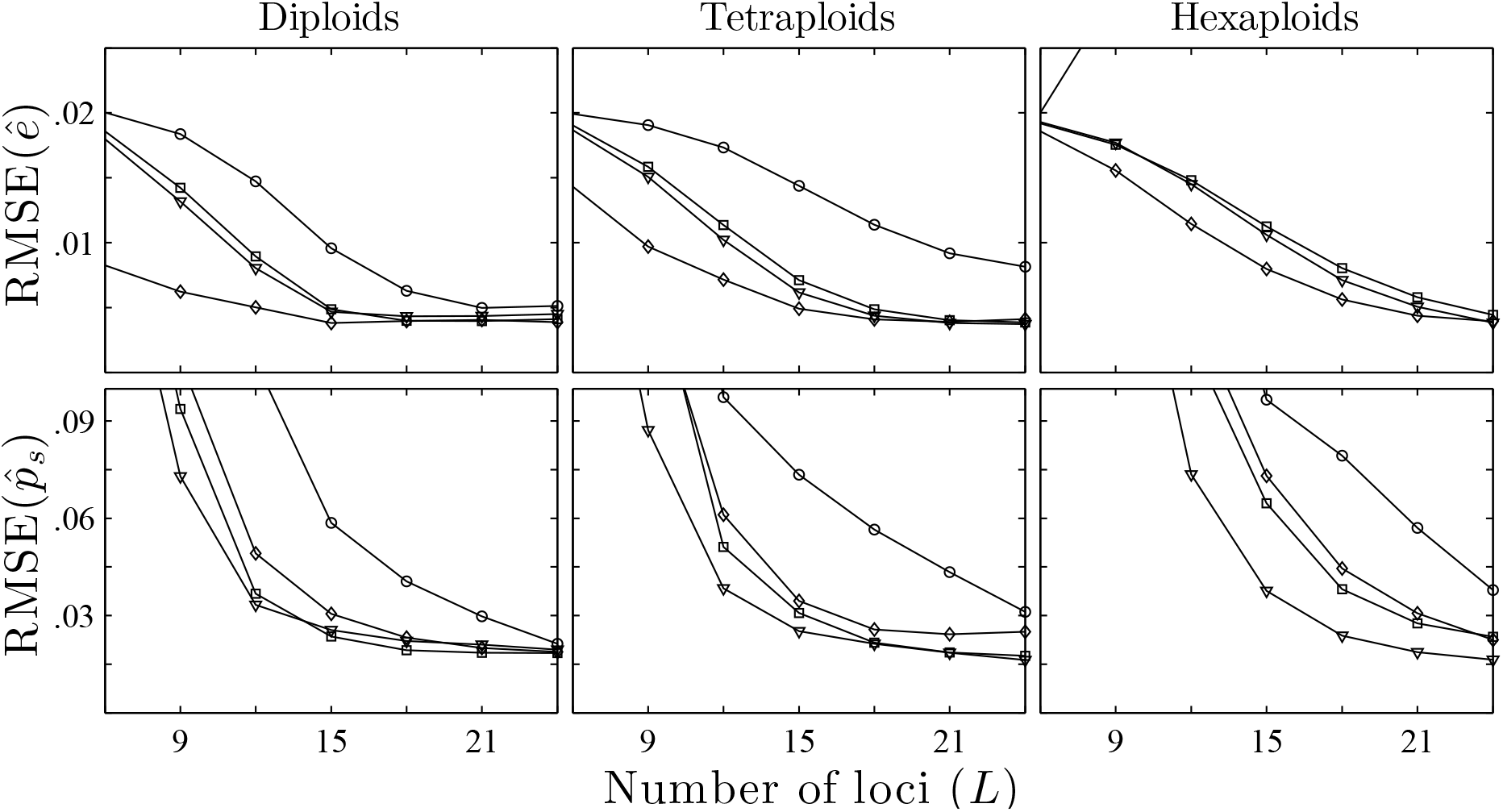
The RMSE of the estimated genotyping error rate *ê* or the estimated sample rate 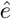 as a function of the number of loci *L* at *e* = 0.02 and *p_s_* = 0.8. The remaining are as for Figure 4.

